# Molecular simulations of enzymatic phosphorylation of disordered proteins and their condensates

**DOI:** 10.1101/2024.08.15.607948

**Authors:** Emanuele Zippo, Dorothee Dormann, Thomas Speck, Lukas S. Stelzl

## Abstract

Understanding the condensation and aggregation of intrinsically disordered proteins in a non-equilibrium environment is crucial for unraveling many biological processes. Active enzymes catalyse many processes by consuming chemical fuels such as ATP. Enzymes called kinases phosphorylate disordered regions of proteins and thus profoundly affect their properties and interactions. Protein phosphorylation is implicated in neurodegenerative diseases and may modulate pathogenesis. However, how protein sequence and molecular recognition of a disordered protein by kinases determine phosphorylation patterns is not understood. In principle, molecular dynamics simulations hold the promise to resolve how phosphorylation affects disordered proteins and their assemblies. In practice, chemically-detailed simulations of enzymatic reactions and the dynamics of enzymes are highly challenging, in particular it is difficult to verify whether implementations of driven simulations are thermodynamically consistent. We can now address this problem with residue-level coarse-grained molecular dynamics simulations, integrating Metropolis Monte Carlo steps to model chemical reactions. Importantly, we show how to verify by Markov-state modeling that the realisation of a non-equilibrium steady state satisfies local-detailed balance. We investigate TDP-43 phosphorylation by the kinase CK1*δ* in simulations, examining patterns of phosphorylation and assessing its preventive role in chain aggregation, which may be a cytoprotective mechanism in neurodegenerative diseases. We find that the degree of residue phosphorylation is determined by sequence preference and charges, rather than by the position in the chain. The phosphorylation frequency is also affected by the phosphorylation patterns, since the interactions between CK1*δ* and TDP-43 actively change after each reaction. For TDP-43, our simulations show condensates dissolution through phosphorylation with kinases binding to the condensates and phosphorylating TDP-43 in the condensates.

## Introduction

Biological systems operate far from equilibrium[1]. The functionalities of cells and of their organelles and compartments are possible only through a very precise self-organization, driven by a continuous injection of energy from the external environment [2]. In the cell, chemical energy is stored, e.g. in the form of ATP molecules among others[3]. This energy is then used to synthesize and degrade molecules through biological cycles. On time scales shorter than physiological changes, microscopic rates are approximately constant and the system enters a non-equilibrium steady states (NESS)[3–5].

Cellular compartmentalisation underpinning biological function is achieved not only by lipid membranes and organelles surrounded by such membranes, but also by phase separation of proteins, giving rise to biomolecular condensates[6]. Membrane-less compartments of phase-separated proteins can concentrate or exclude molecules and thus organize biochemical processes in time and space, which is analogous to the compartmentalisation provided by lipid membranes. These phase-separated condensates can often act as chemical reactions organizers[7]. However, these condensates of proteins can also age into solid aggregates, which are believed to contribute to neuronal dysfunction and neurodegeneration[8, 9]. As condensates age and become less liquid-like, they frequently lose their biochemical functionalities[10]. Aggregates of intrinsically disordered proteins (IDPs) are often linked to neurodegenerative diseases. Some examples are Tau protein aggregates, associated with Alzheimer’s disease[11], *α*-synucleic aggregates, associated with Parkinson’s disease [12], or TAR DNA-binding protein 43 (TDP-43) aggregates, mostly found in patients with amyotrophic lateral sclerosis (ALS) [13], frontotemporal dementia [14], but also in many patients with Alzheimer’s disease [15].

Proteins within condensates can also undergo chemical reactions themselves [2], driving the system out of equilibrium by dissipating a biochemical fuel, such as ATP. The modification of those proteins by addition of chemical groups, such as phosphate groups, are referred to as post-translational modifications (PTMs). IDRs are not only essential in driving the condensation of proteins, but they are also prime targets of PTMs [16]. PTMs can drastically change the properties of individual proteins[17] and collectively of condensates [18], enhancing[11, 19] or suppressing the condensation and aggregation of IDPs[20, 21]. For instance, it has been shown that chemical reactions can stabilize the size of liquid droplets by suppressing Ostwald ripening [22, 23].

To connect these advances in the understanding of active processes in condensates to the biological roles of proteins, it will be important to elucidate how ATP driven phosphorylation shapes the interactions of intrinsically disordered protein regions (IDRs) of neurodegeneration-linked proteins such as TDP-43. The disordered low-complexity domain (LCD) of TDP-43 is hyper-phosphorylated in disease, and in experiments such a hyper-phosphorylation has been found to suppress TDP-43 condensation and aggregation[24]. Enzymes can add PTMs to IDPs in dilute solution, but enzymatic addition of PTMs may also occur in protein condensates. Recently, it was shown that phase-separated condensates can speed up phosphorylation of Tau protein[25]. Phosphorylation of the TDP-43 C-terminal residues Ser 379, Ser 403, Ser 404, Ser 409, and Ser 410 in patient samples is associated with neurodegenerative disease[26]. TDP-43 is phosphorylated by Casein kinase 1*δ* (CK1*δ*). How the enzymatic phosphorylation of TDP-43 is modulated in dilute solution and how it is affected by protein condensates is not known. The disordered tail of CK1*δ* is auto-inhibitory[27, 28], but how it inhibits TDP-43 phosphorylation is unclear on the molecular scale. IDRs of enzymes have multiple functions, such as auto-inhibition by binding to the active site. IDRs are involved in substrate binding, for instance IDRs can speed up reactions via fly casting effect, where the IDR increases the search volume for the binding of partner proteins[29].

Phase behaviour of intrinsically disordered proteins (IDPs) and the biological functionalities of protein condensates have been studied in the past years using multi-scale molecular dynamics (MD) simulations. Such simulations capture the spontaneous condensation of hundreds or more proteins while maintaining enough chemical detail in the simulations to elucidate sequence-specific interactions of proteins[24, 30]. Comparison to more highly-resolved coarse-grained methods [30–34] or atomistic molecular simulations [35, 36] can then highlight important drivers of protein condensation[24].

However most of these studies assume thermodynamic equilibrium, neglecting the dynamical changes in the properties of individual proteins and protein condensates, as well as the dissipation caused by chemical fluxes. Much progress has already been made in the simulations of mechanically-driven non-equilibrium steady state (NESS), where external mechanical forces give rise to driven dynamics [37]. An important step was the construction of Markov state models to better understand the effects of driving on the molecular scale[38]. Analogously, a biological chemically-driven NESS, such as molecular motors, can be simulated by maintaining a chemical potential difference, i.e. by fixing the ATP to ADP concentration ratio. [4, 39] Chemical reactions could in principle also be modelled via quantum mechanical approaches[40], but these are computationally very demand-ing, which can preclude their application to large-scales dynamics in complex biochemical systems. Recently, exciting progress has been made in integrating chemical reactions in molecular dynamics simulations via neural networks[41]. Even in the case of coarse-grained simulations, chemical reactions have been modelled through the use of reactive beads that can form bonds between molecules [42]. In many cases, one could model chemical reactions in complex system by combining MD with a suitably chosen Monte Carlo (MC) step [39, 43]. Arguably, the absence of a straightforward approach of validating the thermodynamic consistency of simulations of NESS has held back the widespread application of MD/MC approaches to biochemical reactions on the molecular scale.

Here we demonstrate how to validate the thermodynamic consistency of simulations of enzymatic phosphorylation of proteins using TDP-43 LCD and its phosphorylation by CK1*δ* as an example. We do so by constructing a Markov state model (MSM), which is a generally applicable approach. Our coarse-grained simulations of enzymatic phosphorylation of TDP-43, show how the sequence specific interactions of CK1*δ* with TDP-43 LCD affects the phosphorylation frequency of serines residues in the TDP-43 LCD in dilute solution and in condensates. In particular the C-terminal domain is more phosphorylated than the N-terminus, in agreement with experiments. Indeed, multiple serines of TDP-43 LCD have been found phosphorylated in patient samples, in particular in the C-terminal region[44, 45], with Ser 409/Ser 410 phosphorylation being established as a hallmark of TDP-43 pathology in disease [26] and detected, together with Ser 403/Ser 404 and Ser 379, by phospho-specific antibodies[46]. The phosphorylation frequency is also affected by the phosphorylation patterns, since the interactions between CK1*δ* and TDP-43 actively change after each reaction, enhancing further phosphorylations[47]. Moreover we study the role of the CK1*δ* IDR (residues from 295 to 415) in phosphorylating TDP-43 both in condensate and dilute regime. CK1*δ* IDR strongly interacts with TDP-43 LCD, reducing its contacts with active site of the enzyme in dilute regime. In dense regime, the CK1*δ* tail anchors of the enzyme to the droplet surface.

## Results

### Markov-state modeling demonstrates thermodynamic consistency of simulations of chemically-driven dynamics

Molecular dynamics (MD) simulations together with a thermostat holding the temperature fixed can be employed to sample from the canonical equilibrium distribution. However, introducing phosphorylation reactions in MD simulations generally inject energy into the system, thus breaking detailed balance and displacing the system away from thermal equilibrium. We simulate the action of the kinase CK1*δ* (truncated at residue 294 for the purposes of this section) on the substrate protein TDP-43 by combining one-bead-per-residue implicit-solvent MD with MC phosphorylation steps and validate the thermodynamic consistency of our simulations by making use of Markov state models (MSMs). We assume that only the serines (Ser) of TDP-43 LCD can be phosphorylated into phospho-serines (pSer). The phosphorylation reaction is the following:

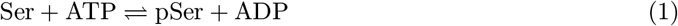

Whenever a Ser (or pSer) is in contact with the active site of the kinase, we try to swap it with a pSer (or the opposite) with acceptance probability given by

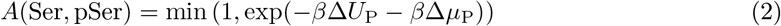

where *β* = 1*/*(*k*_*B*_*T*), Δ*U*_P_ is the difference between the potential energy of the configuration with pSer and the one of the configuration with Ser, and Δ*µ*_P_ is the chemical potential difference between the ATP and ADP molecules involved in the phosphorylation reaction (Eq. 1). ATP and ADP are modelled implicitly and are not explicitly simulated, with concentrations kept fixed and fully characterized through the choice of Δ*µ*_P_. Indeed biological reactions, such as the phosphorylation reaction, in living cells happen in open systems in which the concentrations of substrates, products and the chemical fuel are kept approximately constant over relevant timescales by e.g. metabolic processes. A non-zero value of Δ*µ*_P_ biases the chemical reaction, pushing the simulation away from thermodynamic equilibrium.

The first step is to validate the thermodynamic consistency of our simulations by showing that the energy gain in a phosphorylation cycle (referred to as Δ*µ*_cycle_ in the following) is equal to the chemical potential difference Δ*µ*_P_ in the phosphorylation step. In order to compute Δ*µ*_cycle_, we employ a discretization of the MD trajectory in a MSM. The simplest example of a phosphorylation cycle that we can build is a system with one enzyme and one substrate protein in which only one residue is reactive. In order to get complete phosphorylation cycles, we assume the exchange between TDP-43 and phosphorylated TDP-43 happens when substrate and enzyme are far away from each other without chemical driving and with equilibrium concentrations, through another MC step (Methods). This naturally happens in cells through the action of phosphatases that can catalyze a dephosphorylation reaction.

To gain insights into the effects of including phosphorylation through Eq. 2, we build an MSM from simulated MD trajectories. Firstly we distinguish between bound and unbound state using a neural network called VAMPnet [48]. VAMPnet is able to map molecular coordinates to Markov states through a score function called VAMP-2 score based on the Koopman’s theory. Finding the transformation of the input variables that maximizes the VAMP-2 score is equivalent to optimizing the Markovianity of the output states. In this way we can easily distinguish between the two slowest processes, binding and unbinding, without arbitrarily choosing an a priori criterion of contact. As input for the neural network, we use the 154 distances between each residue of TDP-43 LCD and the active site of CK1*δ*, while as output we ask for 2 states (ideally bound and unbound). We then filter spurious transitions using transition-based state assignment[49]. As an example, we show in Fig. 2a the trajectory of the distance between Ser 403 (the reactive residue) of TDP-43 LCD and the active site of CK1*δ* for the simulation at Δ*µ*_P_ = − 5 kJ/mol (SI Movie 1). We can see that the two states predicted by the neural network comprise bound configurations (when the distance between Ser and CK1*δ* active site is smaller) and unbound configurations (when the distance is larger).

By distinguishing between Ser and pSer along the trajectory, we coarse-grain the system dynamics into the 4 states sketched in Fig. 1. Assuming that our system is a NESS, we can then compute the time-independent transition probabilities *T*_*ij*_(*τ*) from state *i* to state *j* using the non-reversible Maximum Likelihood estimator [50, 51]. We report in Fig. 2b the resulting MSM discretized trajectory referred to the simulation in panel A. Complete cycles 1 →2 →3 →4 →1 are highlighted in red. Every step of the Markov chain corresponds to 10^4^ MD steps, or 0.1 ns in simulation time. For all our simulations, we choose a lag time *τ* = 10 Markov chain steps (Methods). We show in Fig. 2c the implied timescales for the example case of reactive Ser 403 and Δ*µ*_P_ = − 5 kJ/mol. In the end, we estimate the goodness of the MSM by looking at the Chapman-Kolmogorov test (CK test) [52, 53]. In all the validation simulations, the CK tests suggest good agreement between model and prediction for a wide range of lag times, as shown in Fig. 2d for the example case of reactive Ser 403 and Δ*µ*_in_ = − 5 kJ/mol (SI Table S1, Table S2, Fig. S1 for complete data, Methods).

**Figure 1:**
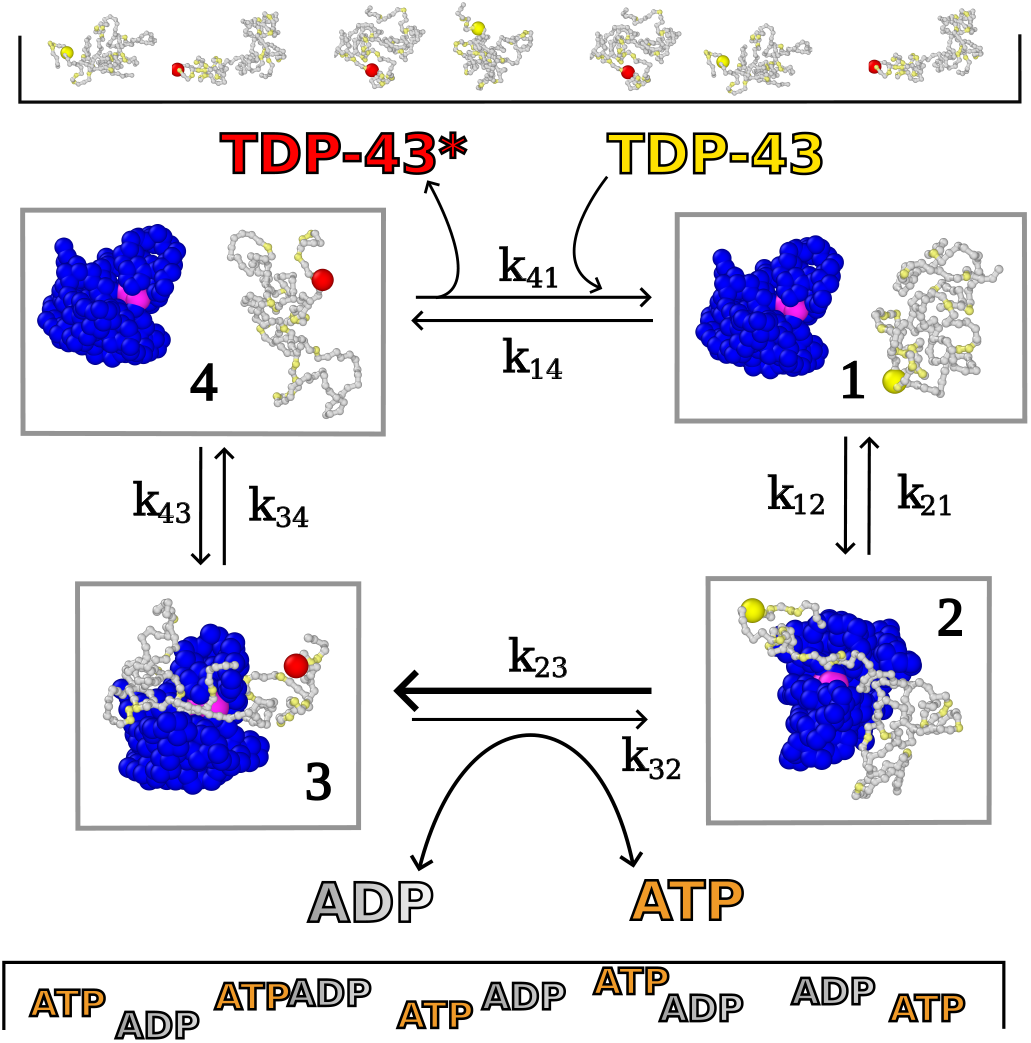
Enzymatic phosphorylation cycle driven by the consumption of the chemical fuel ATP. In state **1** TDP-43 (grey) is unphosphorylated and is not bound to the kinase CK1*δ* (blue, active site in pink). In state **2** TDP-43 binds to CK1*δ*. In state **3** the reactive serine is phosphorylated by kinase, converting one ATP into one ADP. In state **4** phosphorylated TDP-43 dissociates from CK1*δ*. Phosphorylated and unphosphorylated TDP-43 are supplied through reservoirs and we consider exchanges between these reservoirs and our simulation box. Serines are colored in yellow, while phospho-serines in red.

If our system is a NESS, the local detailed balance condition must be satisfied [3, 5]:

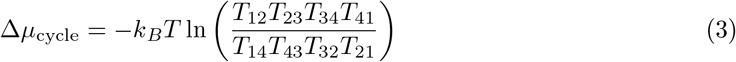

where Δ*µ*_cycle_ is the energy injected into the system in one forward cycle 1 → 2 → 3 → 4 → 1. It is interesting to observe that the logarithm contains the ratio between the forward and backward transition probabilities. In many formulations, Eq. 3 is via rate coefficients rather than transition probabilities. For the short lag times considered here, we can estimate a rate matrix from the transition probability matrix and find virtually indistinguishable results for Δ*µ*_cycle_ (SI Text). Since the transitions 1 ⇌ 2, 3 ⇌ 4 (the binding/unbinding of the enzyme with TDP-43 or phosphorylated TDP-43) and 4 ⇌ 1 (the reservoir exchange step) satisfy detailed balance, while the phosphorylation reaction 2 ⇌ 3 breaks detailed balance injecting into the system an amount of energy equal to Δ*µ*_P_, we expect Δ*µ*_cycle_ to be equal to Δ*µ*_P_ (SI Text). We compute the estimated energy gain Δ*µ*_cycle_ from the transition probabilities *T*_*ij*_ and plot them against the parameter Δ*µ*_P_ of the phosphorylation step for different reactive Ser and Δ*µ*_P_. Encouragingly, for all the six different phosphorylation sites, the chemical potential computed from Eq. 3 matches the applied chemical potential Δ*µ*_P_ (Fig. 2e).

**Figure 2:**
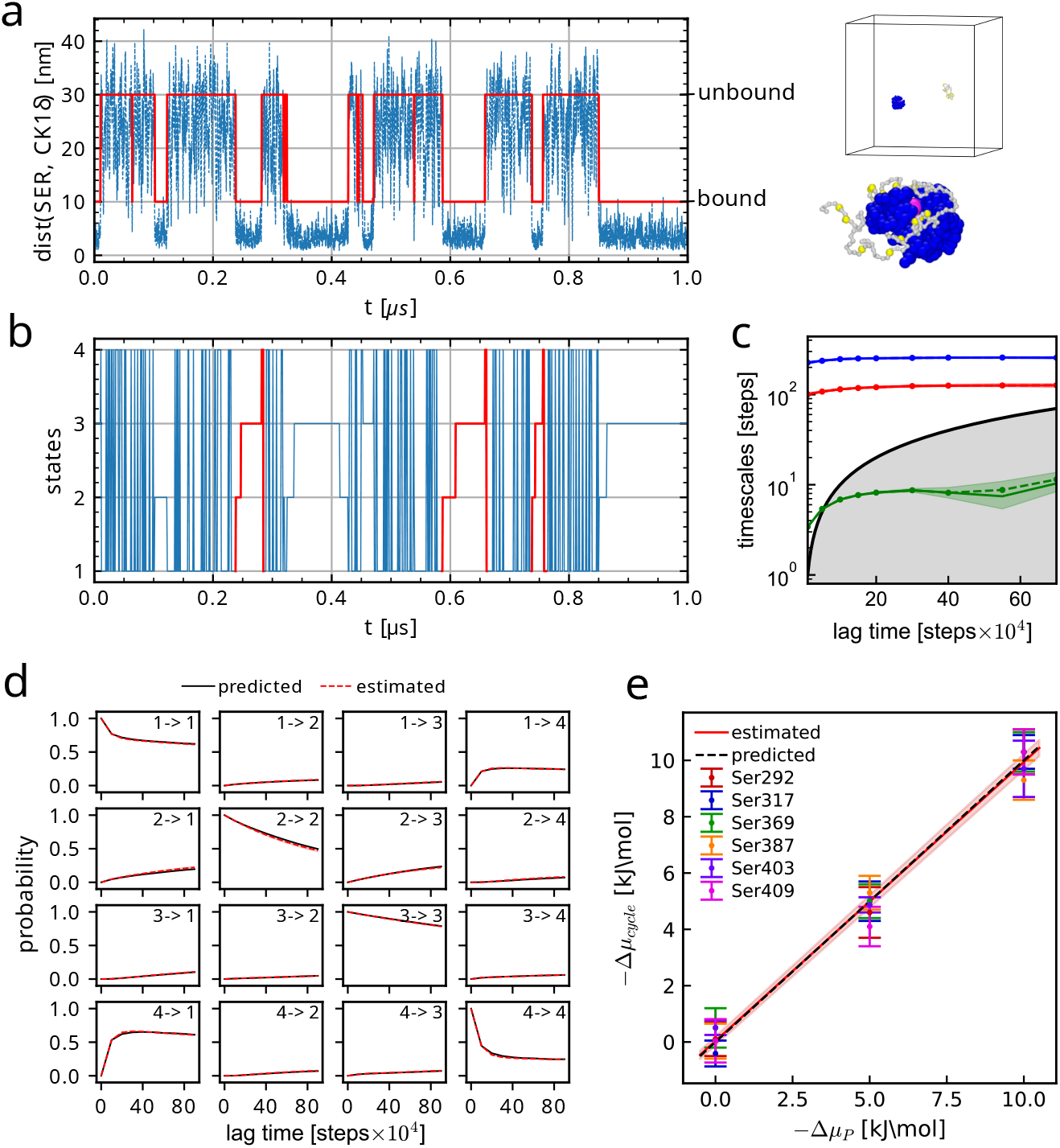
Thermodynamic consistency of simulations with phosphorylation step using MSMs. (**a**) Example of trajectory of the distance between Ser 403 of TDP-43 LCD and the active site of CK1*δ* for the simulation at Δ*µ*_P_ = − 5 kJ/mol. In red the two output states of the neural network (bound and unbound) with respective illustrative examples on the right. (**b**) Example of discretized 4-state MSM trajectory related to the trajectory in **a**, we highlight complete cycles in red. (**c**) Example of implied timescales from the 4-state MSM trajectory shown in **b**; they remain stable for *τ* ≳ 10^5^ MD steps (10 Markov chain steps). (**d**) Example of Chapman-Kolmogorov test from the 4-state MSM trajectory shown in **b**; prediction and model are in agreement. (**e**) We plot Δ*µ*_cycle_ vs Δ*µ*_P_; for all the six different phosphorylation sites, the chemical potential computed from Eq. 3 matches the applied chemical potential Δ*µ*_P_. Errorbars on Δ*µ*_cycle_ are obtained via bootstrapping of the total simulation trajectory collected.

We repeated the estimate of Δ*µ*_cycle_ using a 3-states MSM, in which the unbound states 1 and 4 are merged into the new state 1. The results are in agreement with the 4-states MSM (SI Table S1, SI Fig. S1). Indeed, the transition between state 1 and 4 has a very high rate and can be associated with the smallest implied timescale, that is lower than the lag time for *τ* = 10 Markov chain steps or larger.

We also checked the reliability of VAMPnet by using considerably more input distances (4620 distances) and a different architecture for the case of reactive Ser 403 and Δ*µ*_P_ = − 5 kJ/mol (Methods). The estimated Δ*µ*_cycle_ with the new version of VAMPnet is Δ*µ*_cycle_ = 4.7 ± 0.6 kJ/mol (implied timescales and CK test in SI Fig. S2).

### Phosphorylation preferences are determined by sequence-specific interactions

Having established a model of chemically-driven dynamics, we investigate how sequence context determines the phosphorylation of the disordered protein TDP-43 LCD by the enzyme CK1*δ*, so that we can begin to rationalize sequence-specificity of TDP-43 phosphorylation in experiments[24] and why C-terminal Ser residues such as Ser 410 are frequently found to be phosphorylated in experiments[24, 26, 44]. In our simulations, we follow directly the dynamics of TDP-43 LCD and CK1*δ* folded domain (truncated at residue 294) on the single molecule level (Fig. 3a). We run 100 simulations of TDP-43 LCD in presence of CK1*δ* and at physiological ATP/ADP ratio (Δ*µ*_P_ = − 48 kJ/mol), which mimics in vitro kinase assays. In the simulations, unphosphorylated TDP-43 LCD will eventually encounter CK1*δ* and give rise to different phosphorylation patterns, as shown for an example simulation on Fig. 3a. In this simulation TDP-43 LCD is initially phosphorylated in the C-terminal region. The kinase dissociates after two phosphorylation events and then binds again to the substrate. Multiple Ser residues in the C-terminus of TDP-43 LCD are phosphorylated, including Ser 410, which gets phosphorylated after ten other residues. In our simulations, Ser residues towards the C-terminus of TDP-43 LCD (Ser 369 to Ser 410) are more readily phosphorylated than Ser residues in the N-terminal region of the LCD (Ser 266 to Ser 350), with the phosphorylation rate *r*_P_ on average roughly 3-4 times larger in the C-terminal segment than in the N-terminal segment (Fig. 3b). In mass spectrometry analysis of TDP-43 from ALS patient samples, the phosphorylated sites (the 12 residues Ser 373, Ser 375, Ser 379, Ser 387, Ser 389, Ser 393, Ser 395, Ser 403, Ser 404, Ser 407, Ser 409 and Ser 410 [24, 44] and also Ser 369 [45]) are mostly in the C-terminal region and, interestingly, they are among the ones with largest phosphorylation rate *r*_P_ in our simulations. In particular, Ser 409/Ser 410 phosphorylation has long been established as a hallmark of TDP-43 pathology in disease [26]. This qualitative agreement with simulations tentatively suggests that sequence specific interactions of TDP-43 LCD with the CK1*δ* could explain why these residues are frequently found phosphorylated in experiments and in patient samples.

**Figure 3:**
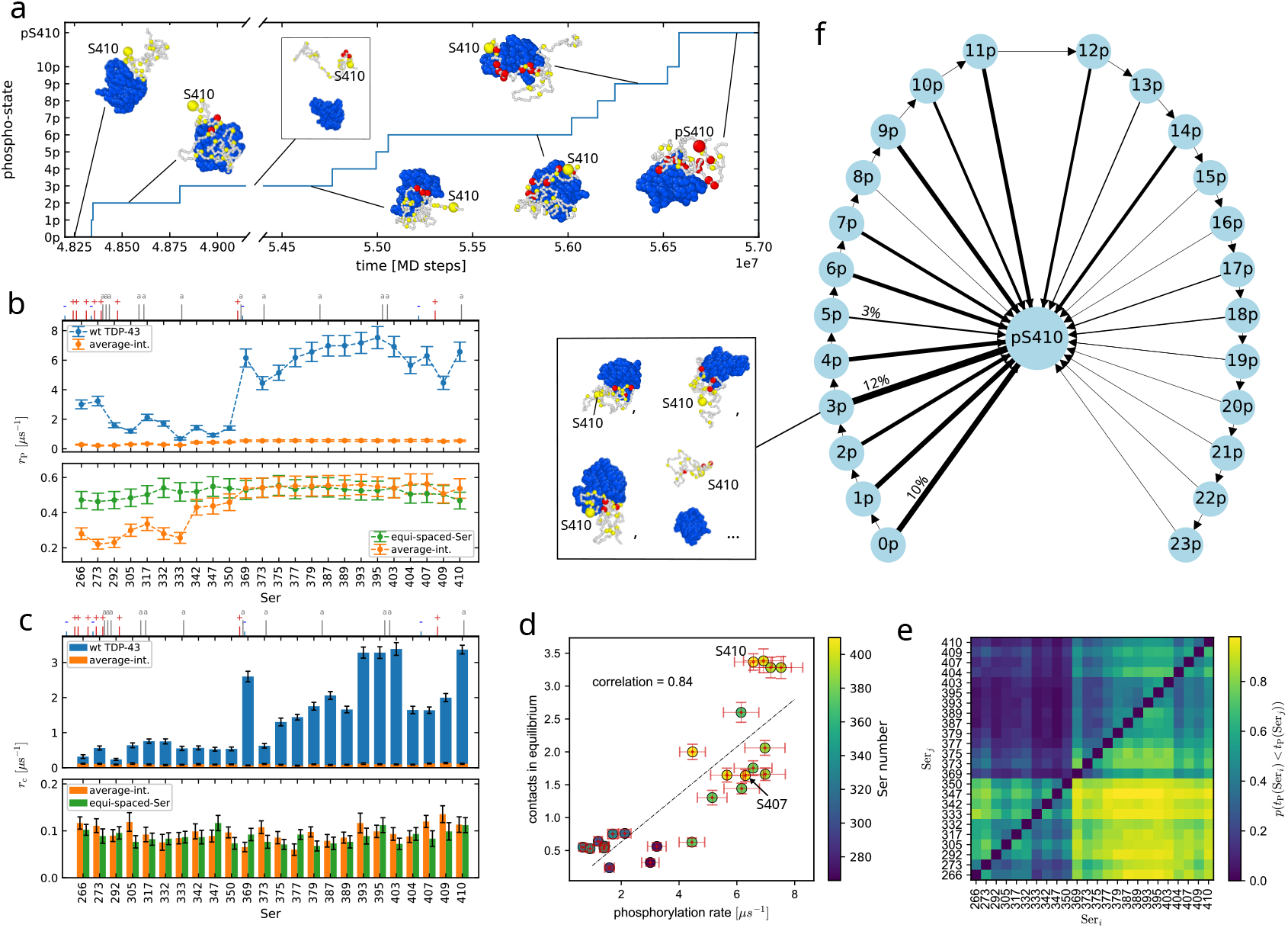
Analyzing the sequence dependence in phosphorylation dynamics of TDP-43. (**a**) Example trajectory discretized in phosphorylation-states relative to Ser 410. *n*p is the state with Ser 410 not phosphorylated and *n* other phosphorylated Ser, pS410 are all the states with phosphorylated Ser 410. (**b**) We compare the phosphorylation rates *r*_P_ for every Ser for the wild type TDP-43 (blue), the averaged-interaction sequence (orange) and the averaged-interaction with equally-spaced Ser (green). The C-terminal is more phosphorylated. The ticks on top show the position of the charged (red ‘+’ positive, blue ‘-’ negative) and aromatic (grey ‘a’) residues. (**c**) Same comparison for the rates of contact *r*_c_ between Ser residues of TDP-43 LCD and the active site of CK1*δ* in equilibrium simulations without phosphorylations. The positive charges in the N-terminal screen the interaction with the enzyme. Contact frequency are constant for the averaged-interaction sequence. (**d**) Correlation plot of contact frequency in equilibrium and phosphorylation rates for the wild type TDP-43. Ser 407 and Ser 410 have similar phosphorylation rate, but the probability of contact of Ser 410 is larger. (**e**) Probability *p*(*t*(Ser_*i*_) *< t*(Ser_*j*_)) of Ser_*i*_ being phosphorylated before Ser_*j*_, data from 100 trajectories. C-terminal residues are much more likely to be phosphorylated before N-terminal residues. (**f**) Phosphorylation pattern representation for Ser 410. The thickness of the arrows represent the percentage of simulations in which Ser 410 was phosphorylated after *n* other Ser residues (e.g. 12% of simulations go from state 3p to pS410). We show in the inset some examples of 3p states.

The differences in the phosphorylation rates can be largely accounted for by how readily Ser residues engage in contacts with the CK1*δ* active site (Fig. 3c), with the phosphorylation rates strongly correlated with a sample Pearson correlation coefficient of 0.84 (Fig. 3d). In order to compare the phosphorylation rates with the frequency of making contacts at equilibrium, we performed MD simulations of the same system without phosphorylation MC steps. To establish to what extent contacts predicts the relative phosphorylation rates, we consider a contact whenever all the three distances to residues Asp 149, Phe 150 and Gly 151 close to the active site are less than 1 nm, in the same way as for the MC phosphorylation step. By contrast, the acceptance probability for the phosphorylation MC step for Ser residues once they are in contact is *>* 0.97 for the entire sequence and the variations in the acceptance probability of the phosphorylation step are not correlated with the variation of the phosphorylation rates (SI Fig. S8). Ser residues in the C-terminal segment of the LCD, including Ser 369, Ser 393, Ser 395, Ser 403, and Ser 410, have the largest tendencies to form contacts, as tracked by *r*_c_, which is the rate at which a residue forms contacts with the CK1*δ* active site (Fig. 3c). At the same time these residues have within the statistical uncertainty the fastest phosphorylation rates of the TDP-43 LCD (Fig. 3b). Ser residues in the N-terminal part of the LCD (Ser 266 to Ser 350) form fewer contacts than serines in the C-terminal segment (Ser 369 to Ser 410), with the exception of Ser 373 in the latter segment, which also forms few contacts with the active site of CK1*δ*. The N-terminus is enriched in charged amino acids (mostly positive) (Fig. 3b and c), which may hinder its binding to the CK1*δ* active site, since the active site features multiple charged residues and is overall positively charged (SI Fig. S7). On the other hand, the C-terminus has more aromatic residues, which increase the attraction through cation-pi and pi-pi interactions [54] (Fig. 3b and c). This difference between the N- and C-terminal segments of the TDP-43 LCD is also apparent on the correlation plot in Fig. 3d, where the N-terminal residues have both low rates and low number of contacts, whereas the C-terminal residues have mostly high phosphorylation rates and many contacts with the active site.

### Dynamics of TDP-43 serine phosphorylation is influenced by preceding phosphorylation events

Although the correlation between the relative rates for CK1*δ* and TDP-43 contact formation and the phosphorylation rates is strong, there are deviations from the this simple relationship (Fig. 3d), which could hint at structural correlations and possible correlations between phosphorylation events. For instance, Ser 410 forms contacts more than two times more readily than Ser 407 but their phosphorylation rates are the same within the statistical uncertainty (Fig. 3b and Fig. 3c). To better understand the underlying correlations, we expanded our analysis of the phosphorylation kinetics. To estimate the phosphorylation rates *r*_P_, we assume that the phosphorylation process is a memory-less process, which follows single-exponential kinetics [55]. In this case, observing a single event is in principle sufficient to estimate the rates of a process. In addition to the number of events one observes, the time spent waiting before an event happens also contributes to the rate estimate. We checked the results by fitting the cumulative histograms of phosphorylation time for each Ser with a simple single-exponential process and an exponential process conditioned to another exponential process (e.g. the binding of TDP-43 to CK1*δ*) (Methods). Most of the times the conditioned exponential process fits perfectly. We found that the rate extrapolations from the two fits are in agreement with the Bayesian estimates (SI Fig. S9). It is interesting to notice that the fastest rate is different for every Ser (Ser 266 with a second rate coeffiecent of 13.5*µ*s^−1^ and Ser 393 of 56*µ*s^−1^) suggesting that the phosphorylation of some serines could involve other processes than the binding to CK1*δ*, e.g. the previous phosphorylation of another Ser. For Ser 410, the two fit extrapolation and the single-exponential fit are in agreement, with differences in the phosphorylation rate of about 2%, while for Ser 403 the conditioned process fit leads to an 8% smaller rate compared to the single-exponential fit. For Ser 407, the conditioned process yields a 10% larger rate. These comparison suggests that the phosphorylation of Ser 403 and Ser 407 could actually follow a more complex process.

We determined the most likely order of phosphorylation to understand correlation between phosphorylation events and differences from what the contact statistics at equilibrium would predict better. In order to study more deeply the phosphorylation pattern of TDP-43, we count for each Ser couple (Ser_*i*_, Ser_*j*_) how many times Ser_*i*_ is phosphorylated before Ser_*j*_ by aggregating data from our 100 trajectories to compute the probability *p*(*t*_P_(Ser_*i*_) *< t*_P_(Ser_*j*_)), where *t*_P_(Ser_*i*_) is the time of phosphorylation for Ser_*i*_ from the start of the simulation. We show *p*(*t*_P_(Ser_*i*_) *< t*_P_(Ser_*j*_)) as a heatmap in Fig. 3e. We see again that, on the single-molecule level, C-terminal residues are typically phosphorylated first. The lower right corner shows that on average C-terminal residues are much more likely to be phosphorylated before N-terminal residues and as a corollary, the upper left sub-matrix shows that C-terminal residues are rarely phosphorylated after N-terminal residues. Instead, looking at the lower left block, we see that Ser 266 and Ser 273 are usually the first phosphorylated in the N-terminal region, while the serines within residues 333 and 350 are the last ones. In the end, by focusing in the C-terminus on the upper right block, we see that the first phosphorylations occur on Ser 369, Ser 393, Ser 395, Ser 403 and Ser 410, followed by Ser between 377 and 389 and Ser 407. In Fig. 3f we aggregate the data from the different trajectories and illustrate the likelihood for Ser 410 of getting phosphorylated after *n* other Ser through the thickness of the arrows. In the figure, the state pS410 includes all the possible configurations in which Ser 410 is phosphorylated, while *n*p are the configurations with *n* pSer different from Ser 410, as shown in the inset for four different examples of state 3p. Very often Ser 410 is among the first three residues to be phosphorylated. Only in a few trajectories, Ser 410 is phosphorylated after nine or eleven other Ser residues are already phosphorylated. Ser 395 shows similar behaviour to Ser 410 (SI Fig. S11). While Ser 403 and Ser 407 are also phosphorylated early on by this analysis, they are less frequently the first Ser residues to be phosphorylated compared to Ser 410 (SI Fig. S11), which is in line with the deviations from single-exponential behaviour (SI Fig. S9. A possible influence of prior phosphorylation can also be detected for Ser 373. Ser 373 forms few contacts but is readily phosphorylated. The phosphorylation rate of Ser 373 is just slightly lower than for Ser 375, which has twice as many contacts. Indeed Ser 369 engages in many more contacts and the rates are just slightly higher than for Ser 373 and Ser 375. Fig. 3e shows that the probability *p*(*t*_P_(Ser_*i*_) *< t*_P_(Ser_*j*_)) of Ser 369 to be phosphorylated before Ser 373 and Ser 375 is approximately 0.8 and 0.7. Fig. S11 (SI) demonstrates that Ser 373 and Ser 375 are phosphorylated when multiple Ser residues are already phosphorylated. Changes in the interaction of CK1*δ* with TDP-43 LCD as more residues are phosphorylated could explain why phosphorylation rates are not fully accounted for by the interaction propensities of the LCD with the active site of the enzyme. By analyzing long equilibrium MD simulations with VAMPnet[48], we find that the phosphorylation facilitates the binding of CK1*δ* to the substrate TDP-43 LCD, with the binding free energy going from 5.0 kJ/mol in the case of wild type TDP-43 LCD to 1.2 kJ/mol for a chain with pSer 395, pSer 403 and pSer 410 (SI Text). As a result, the first phosphorylation events speed up further phosphorylation events, in agreement to what suggested by experiments [47], and we find that, in the simulations of enzymatic phosphorylation of TDP-43, phosphorylated TDP-43 LCD stays attached to CK1*δ* (SI Fig. S3).

### Phosphorylation dynamics is determined by sequence context not relative position to N- and C-termini

The relative position of the Ser residues to the N- and C-termini does not affect the phosphorylation rates. It has been hypothesized that the tendency of C-terminal residues to get phosphorylated could be due to the greater accessibility of residues close to the N- and C-termini of a disordered protein chain [24]. In order to understand whether the phosphorylation pattern is affected by the position of the Ser residues along the TDP-43 LCD chain and not only by the neighboring residues, we repeated the same simulation but replacing all the residues of TDP-43 LCD different from Ser with an averaged interaction strength bead, 1) leaving the serines at their original positions and 2) spreading them equally spaced along the chain. From the contact frequency *r*_c_ in the lower panel in Fig. 3c, we can see that in equilibrium, before any phosphorylation occurs, the probability of contact is uniform along the chain, suggesting that the ends are not a priori more accessible and hence that sequence context and its effects on molecular recognition likely explain the prominence of C-terminal TDP-43 phosphorylation. By looking at the phosphorylation rates *r*_P_ (lower panel in Fig. 3b), we can see that the C-terminal domain is more phosphorylated in the case of simple averaged-interaction beads. We also computed the probability *p*(*t*_P_(Ser_*i*_) *< t*_P_(Ser_*j*_)) for the case of averaged-interaction chain (SI Fig. S10 left), which also demonstrated that the C-terminus is phosphorylated before the N-terminus. This suggests that the negative charges of the pSer also plays a role. These are denser in the C-terminus when TDP-43 gets hyper-phosphorylated. Indeed by distributing the Ser residues at equal distances, phosphorylation rates are constant within the statistical uncertainty. A similar behavior has already been found in experiments for the case of cyclin-dependent kinases phosphorylation of multisite targets[56]. Overall, the phosphorylation rates, as well as the contact frequency in equilibrium, are one order of magnitude smaller in the case of averaged interaction sequence compared to the wild type TDP-43, highlighting once more the importance of the sequence context.

### CK1*δ* binds to TDP-43 condensates and dissolves condensates by hyper-phosphorylation

In our simulations, CK1*δ* folded domain binds to TDP-43 LCD condensates and the LCD condensates dissolve when they are hyper-phosphorylated (Fig. 4a). In cells, TDP-43 often phase-separates into liquid-like droplets which has been linked to the formation of toxic aggregates. Recent experiments have shown that hyper-phosphorylation of TDP-43 LCD can prevent phase separation and aggregation by increasing the solubility of TDP-43 [24]. However it remains unclear whether kinases, such as CK1*δ*, actually bind to TDP-43 condensates, or only phosphorylate TDP-43 in dilute solution. Snapshots from an example simulation with five CK1*δ* enzymes are shown in Fig. 4a (SI Movie 3), with the first snapshot depicting the starting configuration with 200 TDP-43 chains phase-separated in a condensate and the enzymes (blue molecules) randomly placed in the box. After 1 *µ*s of simulation time the enzymes are all attached at the surface of the condensate and they are in the process of phosphorylating several serine residues (pSer in red). Hyper-phosphorylated TDP-43 chains start to disassociate from the condensate, which appears almost entirely dissolved after 5 *µ*s of simulation time in the last snapshot. We report in Fig. 4b the percentage of chains in the condensate (blue, left y-axis) and the percentage of phosphorylated Ser (red, right y-axis) over time from simulations with 1,3 and 5 enzymes, averaged over 4 independent replicas. The percentage of TDP-43 chains in condensate drops over time as the phosphorylation count increases.

**Figure 4:**
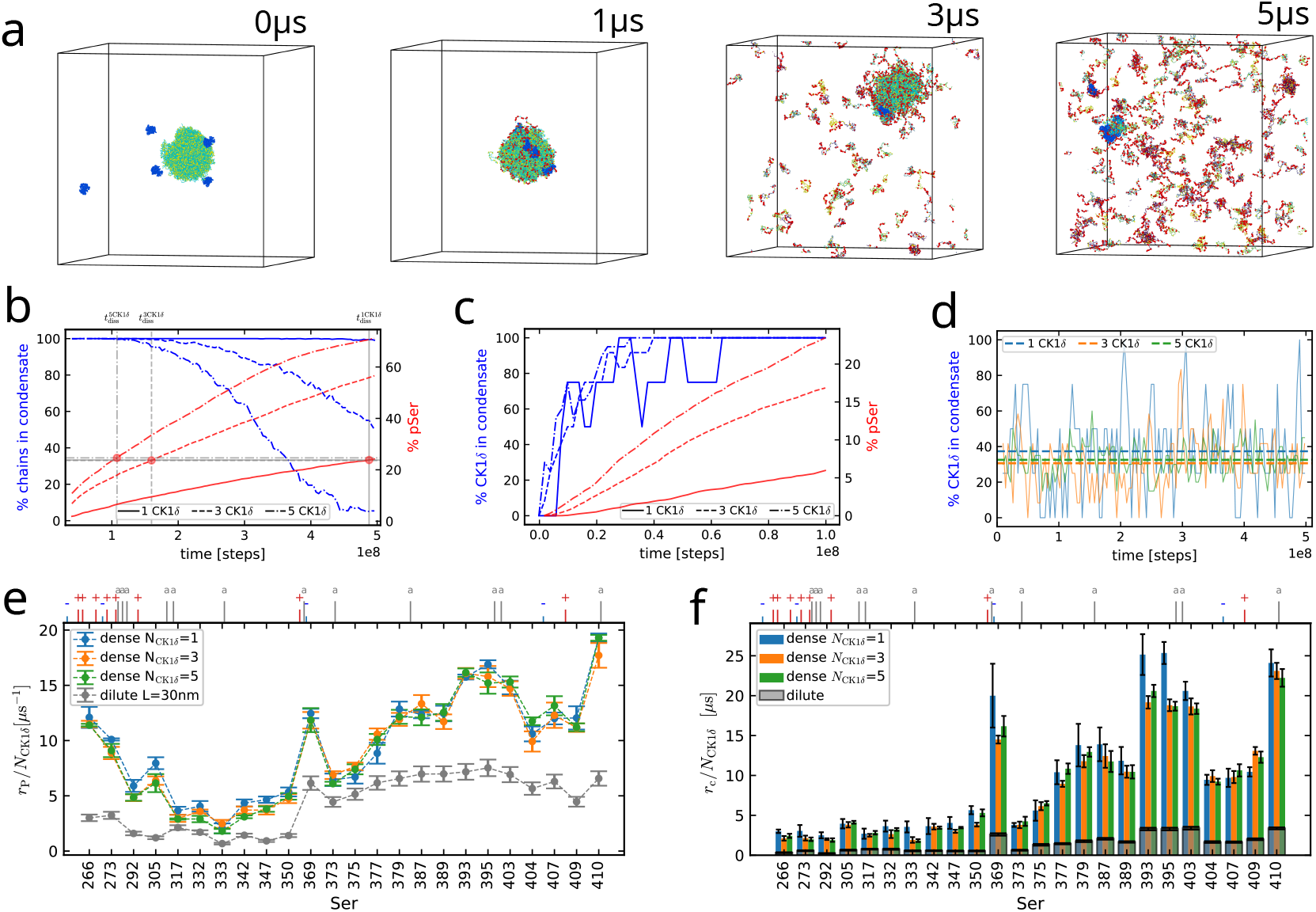
Analyzing the effect of hyper-phosphorylation of a TDP-43 LCD condensate and the interaction of CK1*δ* with the condensate. All the simulations involved in the plots are performed in a cubic box of 100 nm side length with 200 TDP-43 LCD chains. (**a**) Snapshots from simulation with 5 CK1*δ* in cubic box of 100 nm side length at times 0,1,3 and 5 *µ*s showing the dissolution of the TDP-43 condensate. The enzymes are colored in blue and the phospho-serine in red. (**b**) Percentage of TDP-43 chains in the condensate (blue, left y-axis) and percentage of phosphorylated Ser (red, right y-axis) in time for simulations with 1,3 and 5 CK1*δ*. The condensate starts to dissolve after about 24 % pSer. (**c**) Percentage of CK1*δ* attached to the condensate (blue, left y-axis) and percentage of phosphorylated Ser (red, right y-axis) in time for simulations with 1,3 and 5 CK1*δ*. The enzymes remain attached to the condensate after some phosphorylations. (**d**) Percentage of CK1*δ* attached to the condensate in time for equilibrium simulations without phosphorylation with 1,3 and 5 CK1*δ*. In absence of pSer, only about 35% of the enzymes stay attached to the condensate in average. (**e**) Comparison of phosphorylation rates *r*_P_ for every Ser of TDP-43 LCD divided by the number of CK1*δ* chains in dilute regime (grey) and in condensate in presence of 1,3 and 5 enzymes. In dense regime, phosphorylation of the ends of TDP-43 LCD is enhanced. The ticks on top show the position of the charged (red ‘+’ positive, blue ‘-’ negative) and aromatic (grey ‘a’) residues. (**f**) Contact rates *r*_c_ for every Ser of TDP-43 LCD divided by the number of CK1*δ* chains in dilute regime (grey) and in condensate in presence of 1,3 and 5 enzymes at equilibrium.

We computed the size of the condensate using a standard clustering analysis algorithm, thanks to which we were able to distinguish the chains in the larger condensate at every frame (Methods). In every simulation, the condensate starts to lose TDP-43 chains when about 24-25% of Ser are phosphorylated. As a result, the speed of phosphorylation decreases with time, as the TDP-43 chains start to migrate in the dilute regime, far from the action of the enzymes. This effect is particularly evident in the case of 5 CK1*δ* after about 3.5 *µ*s. It is interesting to notice that the speed of phosphorylation decreases slightly with time even before the beginning of the dissolution, with C-terminal Ser being the most affected (SI Fig. S5 upper panel). The slowing down of the phosphorylation rates of the most accessible Ser suggests a possible saturation effect. Moreover, we notice that, at least after about 5% of Ser are phosphorylated, most of the TDP-43 chains feature only 1 or 2 phosphates, with a small minority of chains being hyper-phosphorylated (SI Fig. S6 grey), supporting the idea of an early saturation of the most accessible phosphosites.

TDP-43 phosphorylation facilitates CK1*δ* binding to the condensate, compared to unphoshorylated condensates at equilibrium. In Fig. 4c we show the percentage of enzymes attached to the condensate in time compared to the percentage of phosphorylated Ser, averaged again over 4 replicas. With increasing phosphorylations, CK1*δ* binds more stably to the condensate, suggesting that the negative charges of the pSer residues enhance the binding to the enzyme positively charged residues. In equilibrium simulations without phosphorylation only about 35 % of enzymes are attached to the droplet in average (Fig. 4d).

In our simulations, the protein sequence context determines how much a given Ser residue is phosphorylated in the condensates. We compute the phosphorylation rates *r*_P_ for each Ser of TDP-43 LCD from the counts of phosphorylations and we compare them with the single-chain simulations results. For this computation, we use only the part of the simulations before the start of the condensate dissolution. We can see from Fig. 4e that the phosphorylation rates scale proportionally to the number of enzymes in the box. Moreover, the phosphorylation of the N- and C-terminal serines (namely Ser 266 and Ser 410) is enhanced, as well as for Ser 393, Ser 395 and Ser 403, compared to the single-chain case (see correlation plot in SI Fig. S13 left).

The probability of contact with the active site of CK1*δ* in the condensate for every Ser of TDP-43 LCD differs from the single-chain case roughly by a factor 6. The C-terminus is more accessible, in particular Ser 369, Ser 393, Ser 395, Ser 403 and Ser 410, as shown in Fig. 4f, similar to what occurs in dilute regime. For the dense phase, the phosphorylation rates seem very well correlated to contact rates in equilibrium for the C-terminal serines (sample Pearson correlation 0.91), while the end of the N-terminus is more phosphorylated compared to what one would expect based on the contact statistics from equilibrium simulations (SI Fig. S14 left).

### The role of CK1*δ* disordered domain in the phosphorylation of TDP-43 in dilute solution and condensates

In simulations with full-length CK1*δ*, we find that the disordered region of CK1*δ* (residues 295 to 415) slows down the phosphorylation of TDP-43 in accordance with experiments. In experiments truncated CK1*δ* is more active than the full-length enzyme[27]. First, we run simulations in dilute solution. In our simulations CK1*δ* is unphosphorylated and we do not allow possible autophosphorylations of the CK1*δ* IDR[28] (Fig. 5a, SI Movie 4). With the full-length enzyme, phosphorylation is more restricted to a few residues in the C-terminal region and N-terminal serines are almost never phosphorylated (Fig. 5c). This is due to a reduction of the contacts between TDP-43 residues and the active site of CK1*δ*, located in the folded domain of the enzyme. Indeed, in Fig. 5d we can see that also the rates of contact in equilibrium without pSer *r*_c_ are reduced by one order of magnitude. However, the Ser residues from residue 379 to 410 (10 of the 12 Ser residues found phosphorylated in patient samples) have at least three times the phosphorylation rate of the N-terminus also in the case of full-length CK1*δ*, while Ser 373 and Ser 375 are only slightly more phosphorylated than the N-terminal serines. The rate of active-site contact formation and the phosphorylation rates are more correlated in this case than for the folded domain alone (Fig.S12). However Ser 404, Ser 407 and Ser 409 have similar contacts in equilibrium compared to the N-terminal serines, but higher rate of phosphorylation. As neigboring Ser residues become phosphorylated, binding to the active site, and thus phosphorylation, is likely enhanced.

**Figure 5:**
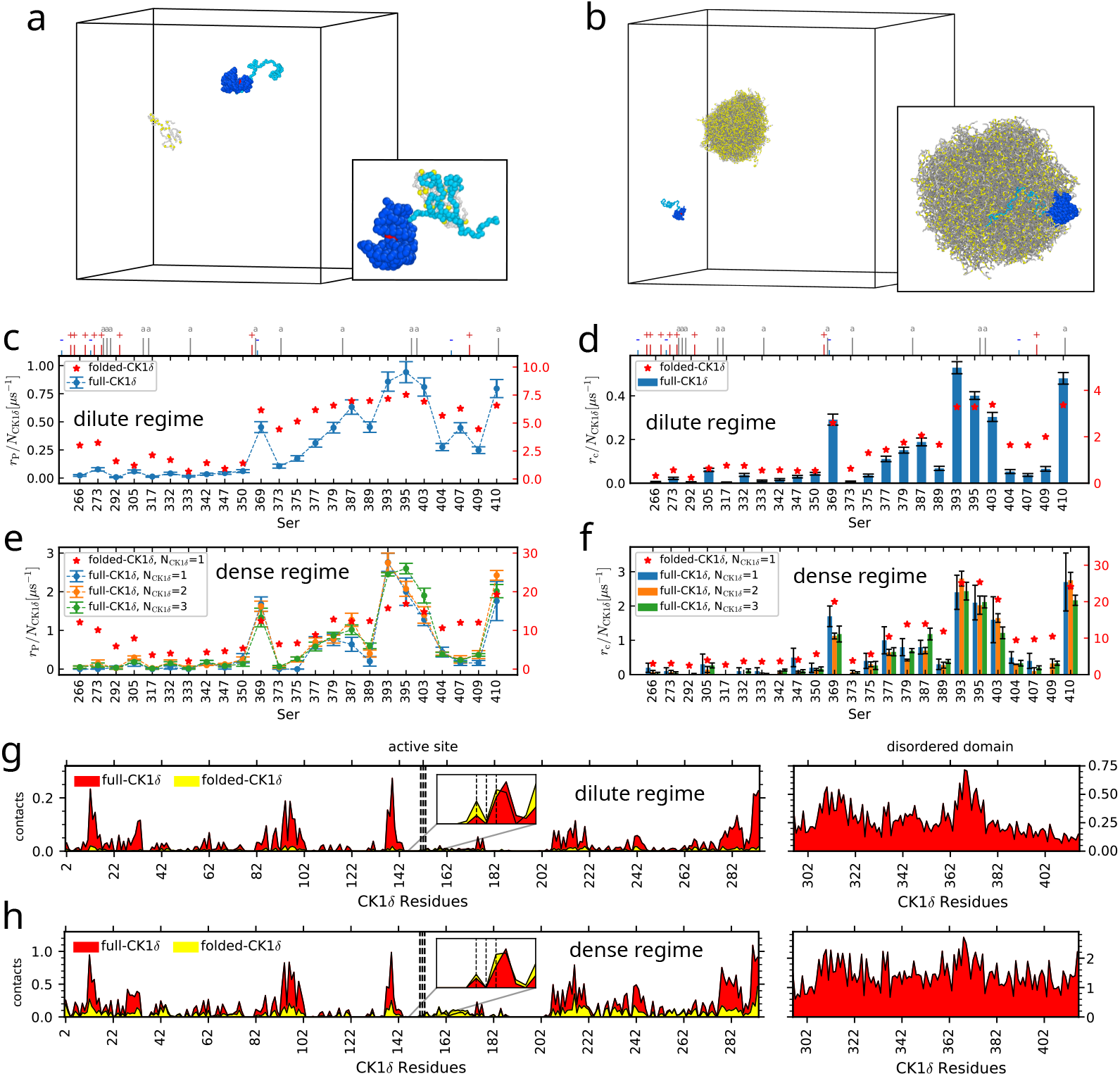
Assessing the role of CK1*δ* disordered domain in phosphorylating TDP-43 LCD both in dilute and dense regime. (**a**) Example of simulation setup of full-length CK1*δ* (blue) and TDP-43 LCD (grey, Ser in yellow) in dilute regime. The inset shows TDP-43 interacting with CK1*δ* IDR (light blue, active site in red). (**b**) Example of simulation setup of full-length CK1*δ* and condensate of 200 TDP-43 chains. The inset shows CK1*δ* IDR (light blue) anchoring the folded domain (dark blue) to the surface of the condensate. (**c-d**) Phosphorylation rates *r*_P_ (c) and contact frequency in equilibrium *r*_c_ (d) for every Ser of TDP-43 LCD in presence of full-length CK1*δ* in dilute regime. The ticks on top show the position of the charged (red ‘+’ positive, blue ‘-’ negative) and aromatic (grey ‘a’) residues. Results from simulations without tail are reported in red on the right y-axis. (**e-f**) Phosphorylation rates *r*_P_ (e) and contact frequency in equilibrium *r*_c_ (f) for every Ser of TDP-43 LCD in presence of full-length CK1*δ* in condensate. Results from simulations without tail are reported in red on the right y-axis. (**g-h**) Contact rates for every residue of full-length CK1*δ* in dilute (g) and dense (h) regime at equilibrium. The disordered region (residues 295 to 415) has more contacts. Simulations with full-length CK1*δ* (red) have more contacts than without IDR (yellow), the insets show that contacts of the active site (Asp 149, Phe 150, Gly 151) are comparable.

The disordered tail of CK1*δ* partially auto-inhibits the enzyme not by occluding the active site, but by sequestering the substrate. TDP-43 LCD interacts more strongly with the tail of CK1*δ* than with the folded domain of CK1*δ* (inset in Fig. 5a). Consequently, full-length CK1*δ* binds more strongly to TDP-43 LCD than the folded domain on its own. From equilibrium simulation of the enzyme and TDP-43 LCD as analysed by VAMPnet[48], the binding free energy energy goes from 5 kJ/mol to -4 kJ/mol. The tail sequesters the substrate in the dilute phase allowing less contacts with the active site, that is located on the opposite side of the enzyme surface, and thus resulting in fewer TDP-43 LCD phosphorylation events. We can see in of Fig. 5g that the disordered domain of CK1*δ* (residues from 295 to 415) has more contacts with TDP-43 LCD compared to the folded domain surface residues in equilibrium simulations without pSer. By looking instead to the residues in the folded domain (from 0 to 294), we notice that the full-length CK1*δ* (red) has in general more contacts than the CK1*δ* without tail (yellow), due to the stronger binding of TDP-43 LCD with the CK1*δ* IDR. However, the active site features a comparable amount of contacts in two cases, with residue 149 having even more contacts in the simulations without tail. By contrast, we find in our simulations that the IDR does not inhibit the CK1*δ* by occluding the active site. The disordered tail of CK1*δ* rarely forms close contacts with the active site and any close contacts are lost very rapidly (Fig. S15).

We run simulations with 200 chains of TDP-43 LCD in a cubic box of 100 nm side length, adding 1,2 and 3 chains of full-length CK1*δ* (Fig. 5b, SI Movie 5). In the dense regime there is high amount of chains, so both contacts and phosphorylation counts increase compared to the dilute case (Fig. 5 e and f). Contacts and phosphorylation counts are also highly correlated in this case, since the abundance of chains allows the enzyme to neglect the less accessible phosphosites and phosphorylate the most accessible ones from every chain. This constitutes a disadvantage for Ser 389, Ser 404, Ser 407 and Ser 409 that are less phosphorylated compared to the dilute case. We note that the phosphorylation rates are directly proportional to the number of enzymes acting on the condensate, as well as the contact statistics (SI Fig. S14). By comparing the phosphorylation rates in condensate with the simulations without IDR, we notice that they are in general lower for the full-length case. The tail acts as a filter, allowing only the phosphorylation of Ser 369, Ser 393, Ser 395, Ser 403, Ser 410 and to a lesser extent Ser 377, Ser 379, Ser 387 and Ser 389, apart from some other very rare phosphorylation events. The effect of the tail on the relative phosphorylation rates is even more pronounced than what we observed in the simulations of CK1*δ* and single chains of TDP-43 LCD.When more than 5% of Ser residues in TDP-43 LCD are phosphorylated, most of the TDP-43 chains feature 1 or 2 phosphates. The distribution of the number of phosphorylated Ser residues per TDP-43 LCD chains is even narrower than in the simulations without the CK1*δ* disordered tail (SI Fig. S6 orange). As for the case without tail, the phosphorylation rate of the most accessible phosphosites decreases with time (SI Fig. S5. This saturation effect of the most reactive Ser residues is apparent even before the eventual dissolution of the condensate (SI Fig. S5 lower panel).

The disordered tail of CK1*δ* facilitates the binding of CK1*δ* to TDP-43 condensates. We observe that the tail recruits the condensate and keeps the folded region anchored to its surface, as illustrated in the inset of Fig. 5b. Also in this case the disordered domain of CK1*δ* has more contacts compared to the folded domain surface residues, as shown in Fig. 5h. Thanks to the disordered tail, the enzyme remains bound to the condensate surface even in absence of phosphorylations. As a consequence, the residues of the folded domain of CK1*δ* form more contacts with TDP-43 compared to the case without the tail (Fig. 5h, residues from 0 to 294). However, the number of contacts of TDP-43 with the active site are again comparable in the two cases, explaining why the rates of binding to the active site shown in Fig. 5f are not greater than the ones in Fig. 4f. As for the single-chain simulation, the inaccessibility of the active site seems to be due to its opposite location on the enzyme surface compared to the disordered tail. Despite the stable anchoring to the condensate, the enzyme active site faces outwards, making it less accessible to the TDP-43 serines. Even in this case the auto-inhibitory and self-regulatory effects of the enzyme tail do not seem to be due to an obstruction of the active site, since CK1*δ* IDR interacts strongly with the condensate (SI Fig. S16).

## Discussion

We have demonstrated how Markov-state models enable us to straightforwardly validate molecular simulations of chemically-driven non-equilibrium steady states (NESS). Chemically-driven NESS are essential in cell biology[1]. Cells require the constant turnover of fuels and metabolites to grow and thrive. Chemically-driven NESS are likely also essential in the function of biomolecular condensates in the cells[2].

We envisage that our approach to establish the thermodynamic consistency of simulations and the combination of molecular dynamics and Monte Carlo can be readily applied to more complex systems and simulations of such systems in high resolution [41]. For more complex systems, extracting kinetically meaningful states becomes even more challenging. In this respect advances based on neural networks and Koopman theory are highly encouraging[48, 53, 57, 58].

PTMs such as phosphorylation of proteins are a fundamental regulatory mechanism in cells and with molecular simulations we can start to investigate how protein sequence and structure determine substrate-enzyme interactions and PTM patterns. Our simulations demonstrate that the IDR of CK1*δ* could have important roles in TDP-43 phosphorylation, 1) by facilitating the binding to condensates and 2) by auto-inhibiting the enzyme, which our simulations capture in line with experiments [27, 28]. It is important to note that details of the conformations of proteins will be critical for the molecular recognition of potential phosphorylation sites by kinases and more detailed molecular simulations [59] will be required to fully understand the recognition mechanisms. A more detailed description of conformational flexibility will be particularly important to understand in detail how the disordered tail of CK1*δ* inhibits phosphorylation and whether sequestering of the disordered substrate rather than binding to the active site really underpins auto-inhibition by the CK1*δ* IDR. Overall our simulations point to a potential preference for the C-terminal residues of TDP-43 on account of its sequence. Aggregated TDP-43 in patient samples is frequently phosphorylated at, e.g., Ser 379, Ser 403/Ser 404 and Ser 409/Ser 410 [26, 46], which are among the most phosphorylated residues also in our simulations. Due to the high concentration of substrates in condensates, proteins are readily phosphorylated. The phosphorylation rates for Ser residues are larger for TDP-43 in condensates than in the dilute phase. Interestingly, phosphorylation patterns are overall similar in dilute solution and condensates. While there are differences in the phosphorylation propensities, sequence context still determines which sites can be phosphorylated. One can speculate that differences between phosphorylation in dilute and dense solution could be partly explained by the overall higher phosphorylation level in condensates, which means that some sites will effectively be more readily phosphorylated than in dilute solutions[25], while the kinase may retain sequence-dependent recognition of substrates in the condensates.

## Methods

### Coarse-grained MD simulations

In our work, we simulated TDP-43 LCD and the kinase CK1*δ* using a one-bead-per-residue coarse-grained model called hydrophobicity scale model [31] (HPS model) and a modified version of it, referred to as modified HPS model in the text (SI Text). In these models, the water solvent and the ions concentration are implicit in the pair potential definition. We used the original HPS for the thermodynamic consistency validation simulations, in which we preferred to give priority to the frequency of the binding and phosphorylation events at the expense of having a more realistic force field, in order to get better statistics. For the other simulations, we used the modified HPS model [30] in which cation-pi interactions are enhanced [60] and folded domains interaction are reduced by 30% [61, 62] (SI Text). Simulations were conducted using Langevin dynamics at a temperature of 300 K and friction coefficient of 0.001 ps^−1^ and in a cubic box with periodic boundary conditions of side length of 30 nm for the single TDP-43 chain simulations and 100 nm for the condensate simulations. The simulated TDP-43 LCD includes residues from 261 to 414 of the full-length TDP-43. The folded domain of CK1*δ* (residues from 1 to 294) follows a rigid body dynamics with rotational drag coefficient of 4 ps^−1^ for every axis, the structure is provided by https://alphafold.ebi.ac.uk/entry/P48730.

For the dilute regime, 100 simulations with phosphorylation reaction step and without reservoir exchange step were run, 2 × 10^8^ MD steps long (2 *µ*s in simulation time) for the case with 1 wild type TDP-43 LCD and 1 CK1*δ* folded-domain, 4 × 10^8^ MD steps long (4 *µ*s in simulation time) both for the case with 1 averaged-interaction polymer and 1 CK1*δ* folded-domain and for the case with 1 wild type TDP-43 LCD and 1 full-length CK1*δ*. To characterize the intrinsic affinity of the enzyme for TDP-43 LCD, we repeated the same simulations, but without phosphorylation reactions at thermodynamic equilibrium. We collected in total 450 *µ*s of simulation time for the case of wild type TDP-43 LCD and CK1*δ* folded-domain and 900 *µ*s for the averaged-interaction polymer and for full-length CK1*δ*. The averaged interaction polymer is built by substituting the TDP-43 LCD residues different from Ser with a bead having average TDP-43 LCD mass, size parameter *σ* and hydropathy parameter *λ* (SI Text) 1) leaving the serines at their original positions and 2) spreading them equally spaced along the chain.

We also simulated a condensate of 200 TDP-43 LCD chains. We ran 4 simulations 5 × 10^8^ MD steps long (5 *µ*s in simulation time) with phosphorylation steps without reservoir exchange step (as for the single chain simulations) in presence of 1,3 or 5 CK1*δ* folded-domain chains and of 1,2 or 3 full-length CK1*δ* chains. We repeated the same simulations, but without phosphorylation reactions at thermodynamic equilibrium, collecting a total of 20 *µ*s of simulation time for each case.

All the simulations involved in this paper were performed using the Python package HOOMD-blue version 3.8.1. The code used for the simulations is available at https://github.com/ezippo/hoomd3_phosphorylation. The Ashbaugh-Hatch pair potential for the non-bonded interactions is available at https://github.com/ezippo/ashbaugh_plugin as a HOOMD-blue plugin.

### Phosphorylation reaction through a Monte Carlo step

In addition to the standard MD simulation, we added a Monte Carlo step to mimic the phosphorylation reaction. Every 200 steps of MD simulation, we check if one of the TDP-43 phosphosites is in contact with the active site of CK1*δ*, the area of the enzyme that catalyzes the reaction, identified with the residues Asp149, Phe150 and Gly151. The contact criterion is the following: the three distances between the TDP-43 phosphosite and the residues of the CK1*δ* active site must all be less than 1 nm; in case more than one phosphosite is in contact with the active site at the same time-step, only the closest one is taken into account. When a contact occurs, we try to switch the Ser in contact into pSer (or the opposite) with a Metropolis-like acceptance probability in Eq. 2. The reverse reaction, that is the exchange of pSer with Ser, can also occur with probability *A*(pSer, Ser) = min (1, exp(*β*Δ*U*_P_ + *β*Δ*µ*_P_)), but it is less likely to happen when there is a chemical potential difference favouring the protein phosphorylation. ATP, ADP are modelled implicitly and are not explicitly simulated, with concentrations kept fixed and fully characterized through the choice of Δ*µ*_P_.

The chemical potential difference in a reaction in units of *k*_*B*_*T* is given by the logarithm of the product to substrate concentration ratio. Considering that the ATP concentration in cells is around 1 mM, the concentration of ADP is around 10 *µ*M and fixing a temperature *T* = 300 K for our simulations, we get a chemical potential difference for a phosphorylation reaction Δ*µ*_P_ = *µ*_*ADP*_ − *µ*_*ATP*_ ≃ − 11.5 kcal/mol ≃ −48 kJ/mol (SI Text). Observe that the ATP concentration is two orders of magnitude larger than the ADP concentration, leading to a large negative Δ*µ*_P_ that favors the exchange of Ser into pSer and disfavors the opposite reaction. Moreover we can mimic the ATP to ADP concentration ratio by changing the chemical potential difference in our simulation at fixed temperature. We used Δ*µ*_P_ = 0, −5, −10 kJ/mol for the validation of the thermodynamic consistency simulations and Δ*µ*_P_ = −48 kJ/mol for all the other simulations in dilute regime and condensate.

### Dephosphorylation step

In our validation simulations, we assume the exchange between TDP-43 and phosphorylated TDP-43 happens without chemical driving and with equilibrium concentrations, through another Metropolis-like step (reservoir exchange step). Every 200 MD steps, we check if the distances between the TDP-43 phosphosite and the 3 residues of the CK1*δ* active site is larger than 25 nm (half box side length). In that case, we randomly swap the pSer of the phosphorylated TDP-43 with a Ser (or the opposite) with a Metropolis-like acceptance probability:

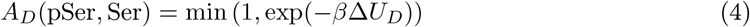

where Δ*U*_*D*_ is again the difference between the potential energy of the configuration with Ser and the one of the configuration with pSer. In this case there is no chemical driving force, the reaction obeys detailed balance and thus it does not inject any additional energy into the system. This exchange step mimics a larger reservoir of TDP-43 and phosphorylated TDP-43 and thus enables us to simulate multiple phosphorylation cycles on the level of a single enzyme and single substrate protein simulation.

### Thermodynamic consistency validation simulations

We simulated the system with one CK1*δ* and one TDP-43 LCD with only one reactive residue. We repatead the simulation for 6 different reactive serines along the TDP-43 LCD, i.e. Ser292, Ser 317, Ser 369, Ser 387, Ser 403, Ser 409, and Δ*µ*_P_ = −0, 5, − 10 kJ/mol. Simulations were conducted for 20 *µ*s in a cubic box of 50 nm side length with periodic boundaries using HPS model force field. In order to get better statistics, we took a 100*µ*s long trajectory for Ser 403 and Δ*µ*_P_ = 0, −5 kJ/mol. We used these longer trajectories for the estimates of Δ*µ*_cycle_ with different lag times and with the version of VAMPnet with more input distances. Errorbars on Δ*µ*_cycle_ were obtained via bootstrapping of the total simulation trajectory collected.

### VAMPnet architecture and training

For the bound state recognition, we performed a nonlinear dimension reduction using a neural network with two identical lobes, following the VAMPnet architecture and the hyper-parameter optimization used by Mardt et al. [48]. Each lobe is composed by an input layer with 154 nodes, one for each residue of TDP-43 LCD, one hidden layer with 30 nodes that employs exponential linear units (ELU) and an output layer with 2 nodes, ideally bound and unbound state, and a final Softmax classifier to obtain probabilities of bound and unbound configurations as output. As input for the neural network, we used the 154 distances between each residue of TDP-43 LCD and the active site of CK1*δ*. We chose a learning rate of 0.5 × 10^−2^ and a batch size of 4 × 10^4^. The neural network was trained for 100 epochs on 90*µs* of equilibrium HPS model[31] simulation with one TDP-43 LCD and one CK1*δ*.

The neural network returns the probability of being in one of the 2 output states (ideally bound and unbound state) for each snapshot of the trajectory. We assigned each snapshot to the state with higher probability, filtering those with a probability between 30% and 70% using transition-based state assignment[49]. In other words, these configurations were assigned based on the state of the previous and following snapshots, in order to filter out spurious transitions.

In order to test the generality of our method, we repeated the bound state recognition with VAMPnet, but using more input nodes. In particular, we used the distances between all the residues of TDP-43 LCD and 30 equally spaced residues of CK1*δ*, resulting in an input layer of 4620 nodes. This time we used 2 hidden layers with 154 and 30 nodes each, and an output layer with 2 nodes. We reduced the learning rate to 0.5 × 10^−3^ and the batch size to 10^4^.

### Implied timescales and Chapman-Kolmogorov test

The choice of the lag time was done by looking at the implied timescales. We can estimate the implied timescales of the Markov model from the eigenvalues of the transition matrix as:

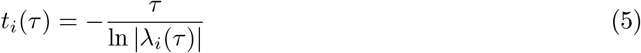

with *λ*_*i*_(*τ*) the eigenvalues of *T*_*ij*_(*τ*). We chose a lag time *τ* such that 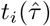 is approximately constant for every 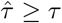. We estimated Δ*µ*_cycle_ for different lag times (1,10,20 Markov chain steps) for the case of reactive Ser 403 and Δ*µ*_P_ = 0, − 5 kJ/mol. For *τ* ≥ 10 Markov chain steps, the estimated Δ*µ*_cycle_ is in agreement with Δ*µ*_P_ (SI Table S2).

We estimated the goodness of the MSM by looking at the Chapman-Kolmogorov test (CK test) [52] [53]. In a Markovian process, the transition matrix satisfies the relation

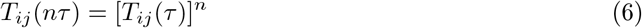

with *n* ≥ 1. In other words, the transition matrix of the model estimated at lag time *nτ* must be equal to the transition matrix to the power *n* of the model estimated at lag time *τ*. The CK test compares *T*_*ij*_(*nτ*) (the estimated transition matrix) and 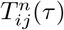 (the predicted transition matrix) for every possible transition *i* ⇌ *j* and different lag times *nτ*.

### Estimate of phosphorylation rates and fit of phosphorylation processes

In all the single TDP-43 LCD chain simulations, we estimated the phosphorylation rates *r*_P_ assuming the phosphorylation process is without memory and thus follows single-exponential kinetics. In all the collected 100 simulations, we had at most one phosphorylation event for each Ser residue, happening at time 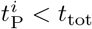 for simulation *i*, with *t*_tot_ the total time of the simulation. In this case, we can use the maximum likelihood estimator for the rate with uniform prior distribution [55]

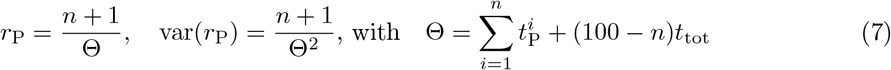

where *n* is the counts of simulations with one phosphorylation event for the Ser took in consideration. Instead, for the simulations in condensate, in which we have multiple TDP-43 chains and thus multiple phosphorylation events for each Ser residue, we computed *r*_P_ as the total count of phosphorylation events in the simulation divided by the total simulation time. In this case, the error on the estimate of the rate is computed as the standard error of the mean from the different replicas. In the same way we also computed all the contacts rates *r*_c_ and their error.

However, since the phosphorylation of a Ser can happen only if TDP-43 is bound to the enzyme, it is more appropriate to take into account the conditional probability of the phosphorylation event given the binding of TDP-43 and CK1*δ* already occurred. Given *p*_*B*_(*t*)*dt* = *r*_B_ exp (−*r*_B_*t*)*dt* the probability of binding between time *t* and *t* + *dt* and *p*_P_(*t*)*dt* = *r*_P_ exp (−*r*_P_*t*)*dt* the probability of having a phosphorylation between time *t* and *t* + *dt*, the conditional probability of having a phosphorylation between time *t* and *t* + *dt* given that TDP-43 is bound to CK1*δ* is

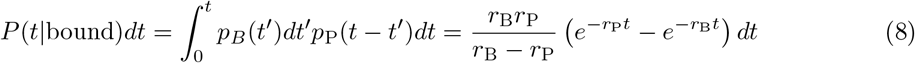

If we call *P*_*c*_(*t < T*) the probability of having a phosphorylation event within time *T* in our simulations, we can write its complementary as:

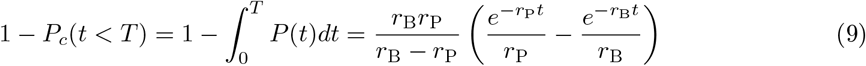

Instead, if we assume that the binding process is much faster than the phosphorylation one (*r*_B_≫ *r*_P_), than we can approximate *P* (*t* | bound) ∼ *p*_P_(*t*) and 1 − *P*_*c*_(*t < T*) ∼ exp (−*r*_P_*t*).

From the 100 simulations used to estimate the phosphorylation rates, we computed the normalized inverse cumulative histogram of the phosphorylation events time *T*, where each time bin gives the phosphorylation counts for that bin plus the counts of all the following bins, divided by the number of simulations. We fitted it with 1 − *P*_*c*_(*t < T*) both for a single-exponential process and a conditioned single-exponential process.

### Condensate identification with clustering analysis

In order to identify the TDP-43 LCD condensate in the trajectory file, we used the DBSCAN (Density-Based Spatial Clustering of Applications with Noise) clustering analysis algorithm[63]. It is an efficient algorithm to identify clusters based on a euclidean distance cut-off *ϵ* and a minimum cluster size parameter *n*_min_. Particles with at least *n*_min_ neighbors within a distance *ϵ* are considered core particles of the cluster. Instead, particles with less than *n*_min_ neighbors are considered non-core particles and they are assigned to a cluster only if at least one of their neighbors is a core particle.

For the estimate of the condensate size and of the percentage of CK1*δ* in contact with the condensate, we used the positions of every bead as input data and we chose the parameters *ϵ* = 1 nm and *n*_min_ = 2. With this choice, every isolated chain is considered as a cluster and two different chains belongs to the same cluster whenever at least one of their particles is in contact (within 1 nm). However, varying *ϵ* between 0.8 nm and 3 nm and *n*_min_ between 2 and 5 does not significantly change the results. We accounted for the periodic boundary conditions by centering the condensate in the box at every frame.

## Acknowledgments

E.Z. was funded by the Deutsche Forschungsgemeinschaft (DFG, German Research Foundation), Project No. 233630050 – TRR 146. L.S.S. thank M^3^ODEL and ReALity (Resilience, Adaptation and Longevity) and Forschungsinitiative des Landes Rheinland-Pfalz for support. This project was funded by the Deutsche Forschungsgemeinschaft (DFG, German Research Foundation) - SFB 1551 – Project No. 464588647. Further, we gratefully acknowledge the computing time granted on the supercomputers Mogon II at Johannes Gutenberg University Mainz, which is a member of the AHRP (Alliance for High Performance Computing in Rhineland Palatinate) and the Gauss Alliance e.V. We thank Alex Holehouse for insightful discussions

## Author contributions

E.Z. ran the simulations, analysed data, interpreted results, wrote the manuscript. D.D. provided important intellectual knowledge and assisted in designing the study, reviewed the manuscript. T.S. conceived the study, provided important intellectual knowledge, reviewed the manuscript. L.S.S. conceived and supervised the study, interpreted results, wrote and reviewed manuscript.

## Supporting information

### Residue-level coarse-grained models with implicit solvent

For our coarse-grained simulations we employed the hydrophobicity scale (HPS) model [31] and a modification of it [30]. The original HPS model was fitted with IDPs data and considers proteins as fully flexible chains. In order to have a more realistic representation of the enzyme CK1*δ*, we decided to employ also a modified version of it that takes into account the presence of folded domains. In both models every residue type is represented with a particle of Lennard-Jones (LJ) size *σ*, charge *q*, mass *m* and hydropathy scale parameter *λ*. For the HPS model, the pair potential has 3 contributions

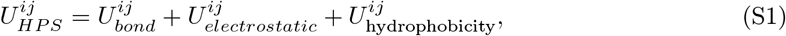

while the modified HPS model has one more contribution to enhance cation-pi interactions:

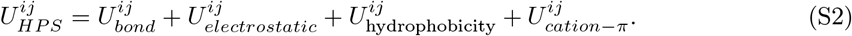

The bonded interactions are described by an harmonic potential

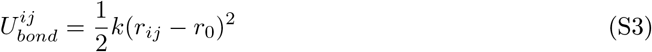

with *r*_*ij*_ the distance between the neighboring residues *i* and *j*, spring constant *k* = 8360kJ*/*(molnm^2^) and equilibrium bond length *r*_0_ = 0.381 nm.

The interactions between non-bonded residues are modeled through the Ashbaugh-Hatch potential

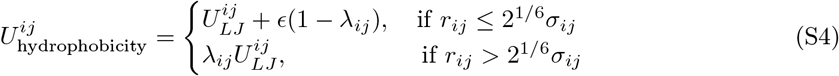

where *σ*_*ij*_ = (*σ*_*i*_ + *σ*_*j*_)*/*2, *λ*_*ij*_ = (*λ*_*i*_ + *λ*_*j*_)*/*2 and 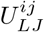 is the standard Lennard-Jones potential

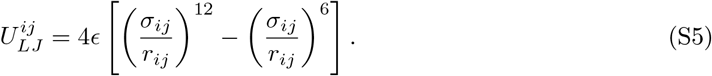

The interaction is truncated at a cutoff distance of 2 nm. The parameter *ϵ* expresses the strength of the Lennard-Jones interaction and it is fixed to *ϵ* = 0.8368 kJ/mol, value fitted with experimental *R*_*g*_ from single IDP chains [31], while the hydrophaty scale parameter *λ*_*ij*_ scales down the inter-action for distances larger than the minimum of 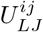 and goes from 0 (fully hydrophilic case, no attraction between residues) to 1 (fully hydrophobic case, the interaction becomes a standard LJ). Phosphorylated Ser residues are modelled as described by Perdikari et al [64].

Charged residues experience also salt-screened electrostatic interactions, which are modeled using a Yukawa/Debye-Hückel potential

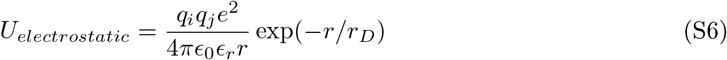

where we used a Debye screening length *r*_*D*_ = 1.0*nm* for an ionic strength of approximately 100 mM and a relative dielectric constant of the water solvent *ϵ*_*r*_ = 80, following the ones of the original HPS model [31]. In this case the cutoff distance is 3.5 nm.

For the modified HPS model, we added another LJ potential only between cation-*π* pairs (Arg/Lys with Phe/Trp/Tyr), as proposed by Das et al. [60]:

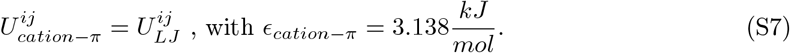

Also in this case the cutoff is 2 nm.

In the modified HPS model, the dynamics of folded domains follows the one of a rigid body. Moreover, the parameter *λ* in 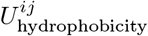 and *ϵ*_*cation*−*π*_ in 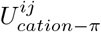 are scaled down by 30% for pair interactions involving residues of the folded domains, as suggested by Krainer et al. [62].

The Ashbaugh-Hatch pair potential for the non-bonded interactions is available at https://github.com/ezippo/ashbaugh_plugin as a HOOMD-blue plugin. The code used for the simulations is available at https://github.com/ezippo/hoomd3_phosphorylation.

### Chemical potential difference in a phosphorylation cycle

The chemical reaction difference in a reaction in units of *k*_*B*_*T* is given by the logarithm of the product to substrate concentration ratio: In the phosphorylation-dephosphorylation cycle, the chemical reactions involved are the two following ones:

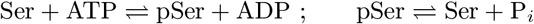

The chemical potential differences for the two reactions are

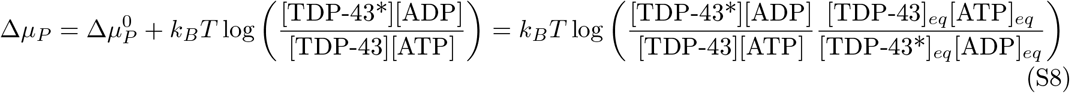

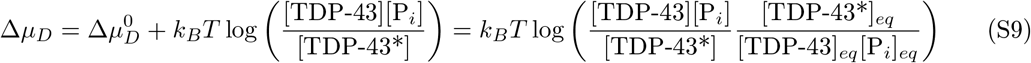

The total amount of chemical driving will be:

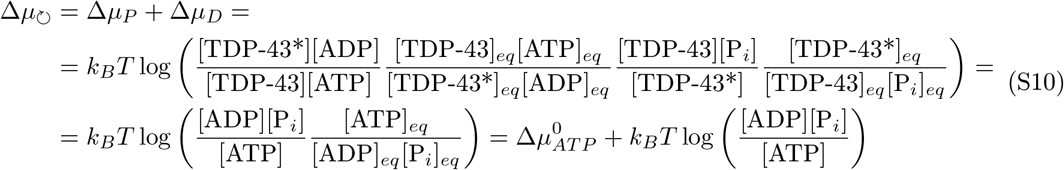

Note that Δ*µ*_⟳_ is equal to the chemical potential difference for the ATP hydrolysis reaction ATP ⇌ ADP + P_*i*_, for which the equilibrium value in standard conditions is 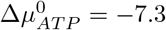 kcal/mol.

### Detailed balance breaking and local detailed balance

In our NESS simulations of a 4-state MSM, we expect to have detailed balance for the transitions 1 ⇌ 2 and 3 ⇌ 4, i.e. the binding/unbinding of the enzyme with TDP-43 or phosphorylated TDP-43, since they are determined by equilibrium MD simulations, but also 4 ⇌ 1, i.e. the reservoir exchange step, that is determined by a Metropolis step without chemical fuel. The detail balance condition is:

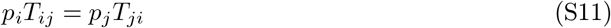

where *T*_*ij*_ is the transition probability from state *i* to *j* and *p*_*i*_ is the stationary probability of being in state *i*. Instead the phosphorylation reaction 2 ⇌ 3 breaks detailed balance injecting into the system an amount of energy equal to Δ*µ*_P_. Despite the Metropolis step is built in such a way to satisfy the detailed balance condition, the Δ*µ*_P_ added in the acceptance ratio breaks detailed balance once the algorithm is coupled to equilibrium MD simulations. In order to show this, let us call ‘A’ a microstate configuration in which we have a contact between the Ser of TDP-43 and the active site of CK1*δ* and ‘B’ the same microstate configuration but soon after the phosphorylation step, with pSer instead of Ser. The probability of being in the microstate ‘A’ or ‘B’ is *p*_*A*_ = exp (−*β*ℋ_*A*_)/Z and *p*_*B*_ = exp (−*β*ℋ_*B*_)/Z, with *β* = 1*/k*_*B*_*T* and Z the canonical partition function, and they are sampled through the MD simulation. Since the velocities in ‘A’ and ‘B’ are the same, the probability ratio will be *p*_*B*_*/p*_*A*_ = exp (−*β*Δ*U*_P_).

The Metropolis acceptance ratio contains also an additional Δ*µ*_P_, leading to the following transition probability for the phosphorylation step:

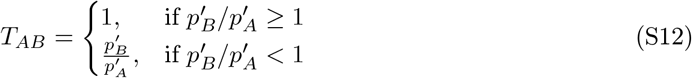

where 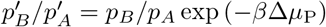. Thus detailed balance is broken for Δ*µ*_P_≠ 0 and we get:

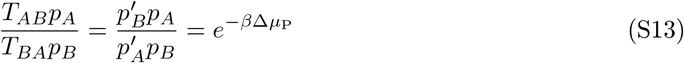

On the other side, if our system is a NESS, i.e. *T*_*ij*_ and *p*_*i*_ are constant in time, for the local detailed balance we should have

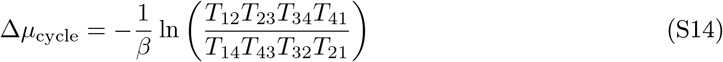

Using Eq. S11 for the couples (*i, j*) = (1, 2), (3, 4), (4, 1), we can simplify Eq. S14 as

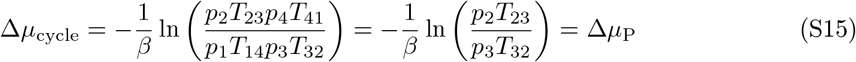

where in the last step we use the local detailed balance condition on the phosphorylation step in Eq. S13.

### Transition probabilities and transition rates

In many formulations, Eq. 3 (or Eq. S14) is expressed in terms of transition rates rather than transition probabilities. The two are related and in our case give the same result for the Δ*µ*_cycle_. We estimated the time-independent transition probability *T*_*ij*_(*τ*), namely the probability of having the system in state *j* at time *t* + *τ* given that it was in state *i* at time *t* (for every *t*), using the non-reversible Maximum Likelihood Estimator[50, 51]:

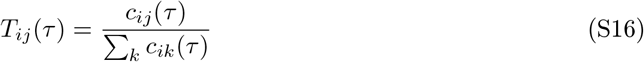

where *c*_*ij*_(*τ*) is the count of transitions from *i* to *j* after a lag time *τ*.

In principle, it could be useful in some cases, e.g. for continuous-time systems or non-linear reaction networks, to express Eq. 3 with transition rates *k*_*ij*_. For a Markov process, the transition probability matrix **T**(*τ*) can be expressed in terms of transition rates matrix **k** as:

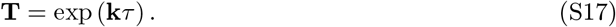

For small *τ* compared to the system timescales, Eq. S17 can be approximated as

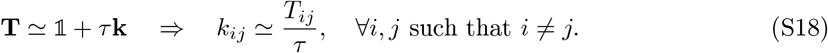

In such case, using *T*_*ij*_ or *k*_*ij*_ to compute Δ*µ*_cycle_ does not change the result, since the factor 1*/τ*would be cancelled out in the ratio in Eq. 3 (or Eq. S14).

As an example, for the simulations with reactive Ser 403 and Δ*µ*_P_ = − 5 kJ/mol, we computed *T*_*ij*_ with lag time *τ* = 10 Markov chain steps (or 1 ns in simulation time). Discretizing the trajectory into 3 Markov states (state 1 with unbound CK1*δ* and TDP-43, state 2 with bound configuration and unphosphorylated Ser 403 and state 3 with bound configuration and phosphorylated Ser 403) leads to 2 implied timescales that are much larger than the the lag time *τ* (Fig. 2, SI Fig. S1). The estimated transition probability matrix is

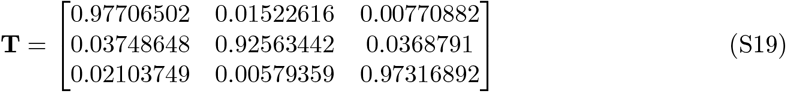

while the transition rates matrix is

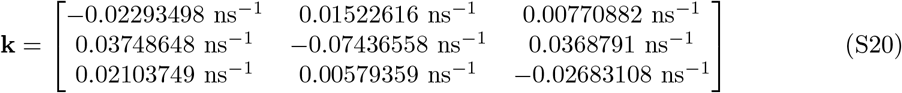

where for the diagonal elements we used the property of the rate matrices *k*_*ii*_ = − Σ_*j≠i*_ *k*_*ij*_.

### Thermodynamic consistency data

**Table S1:** Estimated Δ*µ*_cycle_ reported in Fig. 2E of the main text.

| Ser | $-\Delta\mu_P$ [kJ/mol] | $-\Delta\mu_{\text{cycle}}$ [kJ/mol] | $-\Delta\mu_{\text{cycle},3s}$ [kJ/mol] |
| --- | --- | --- | --- |
| Ser292 | 0 | $0.1 \pm 0.6$ | $0.1 \pm 0.6$ |
| | 5 | $4.6 \pm 0.9$ | $4.6 \pm 0.9$ |
| | 10 | $10.3 \pm 0.8$ | $10.3 \pm 0.9$ |
| Ser317 | 0 | $-0.4 \pm 0.5$ | $-0.5 \pm 0.5$ |
| | 5 | $5.0 \pm 0.7$ | $4.9 \pm 0.7$ |
| | 10 | $10.3 \pm 0.6$ | $10.1 \pm 0.8$ |
| Ser369 | 0 | $0.5 \pm 0.7$ | $0.5 \pm 0.6$ |
| | 5 | $5.0 \pm 0.6$ | $5.1 \pm 0.5$ |
| | 10 | $10.3 \pm 0.7$ | $10.2 \pm 0.5$ |
| Ser387 | 0 | $0.0 \pm 0.6$ | $0.1 \pm 0.4$ |
| | 5 | $5.3 \pm 0.6$ | $5.2 \pm 0.5$ |
| | 10 | $9.3 \pm 0.7$ | $9.1 \pm 0.7$ |
| Ser403 | 0 | $0.51 \pm 0.26$ | $0.52 \pm 0.27$ |
| | 5 | $4.87 \pm 0.27$ | $4.9 \pm 0.3$ |
| | 10 | $9.7 \pm 1.0$ | $9.5 \pm 0.7$ |
| Ser409 | 0 | $0.0 \pm 0.8$ | $0.1 \pm 0.7$ |
| | 5 | $4.1 \pm 0.7$ | $4.1 \pm 0.5$ |
| | 10 | $10.3 \pm 0.8$ | $10.2 \pm 0.8$ |

**Table S2:** Estimated Δ*µ*_cycle_ for Ser 403 and Δ*µ*_P_ = 0, −5 kJ/mol for different lag times. The results for Δ*µ*_cycle_ are in agreement with Δ*µ*_P_ for *τ* ≥ 10 Markov chain steps.

| Ser | $\tau$ [steps] | $-\Delta\mu_P$ [kJ/mol] | $-\Delta\mu_{\text{cycle}}$ [kJ/mol] |
| --- | --- | --- | --- |
| Ser403 | 1 | 0 | $0.45 \pm 0.23$ |
| | 10 | 0 | $0.51 \pm 0.26$ |
| | 20 | 0 | $0.57 \pm 0.27$ |
| Ser403 | 1 | -5 | $4.29 \pm 0.21$ |
| | 10 | -5 | $4.86 \pm 0.27$ |
| | 20 | -5 | $4.99 \pm 0.34$ |

**Figure S1:**
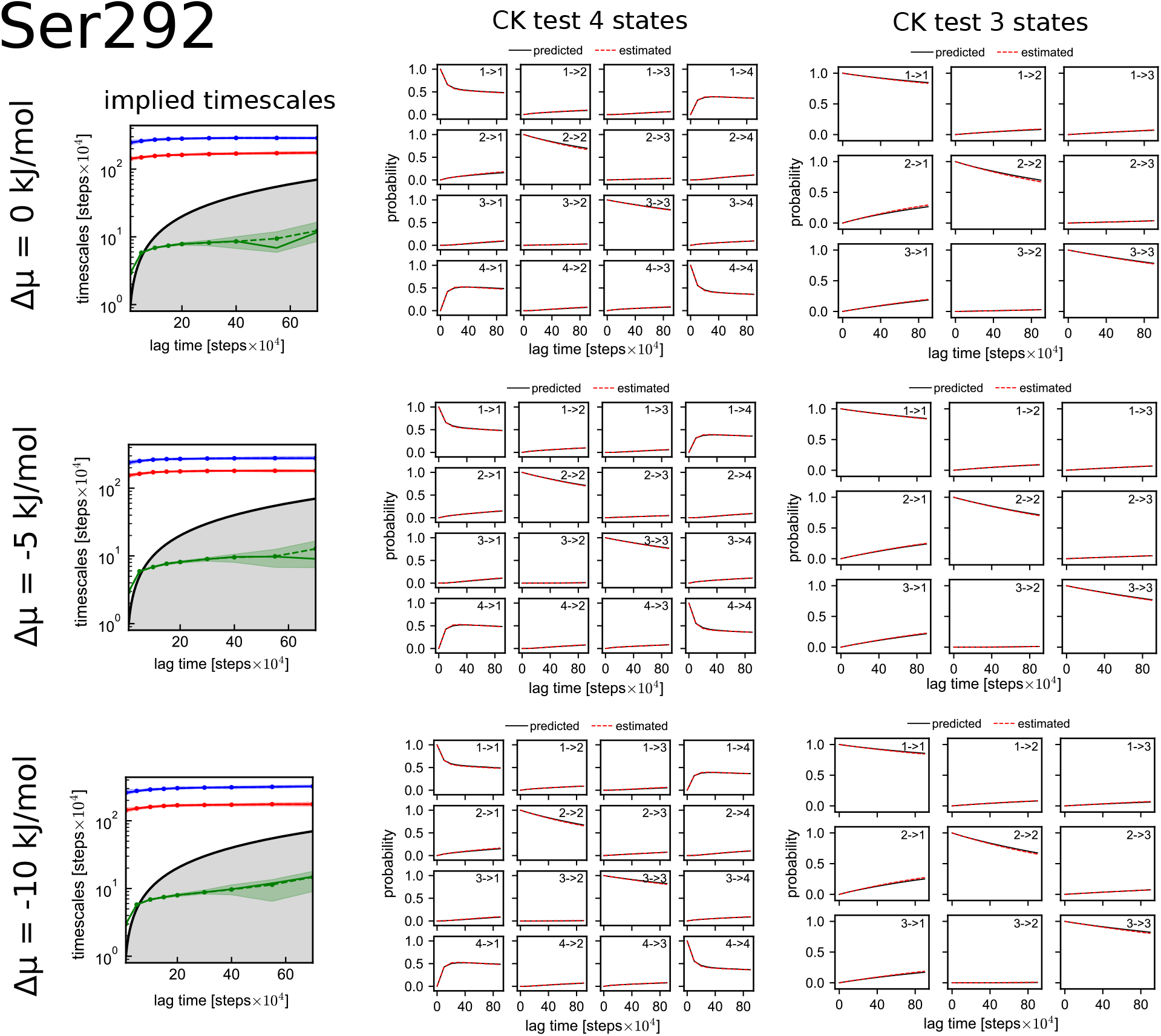
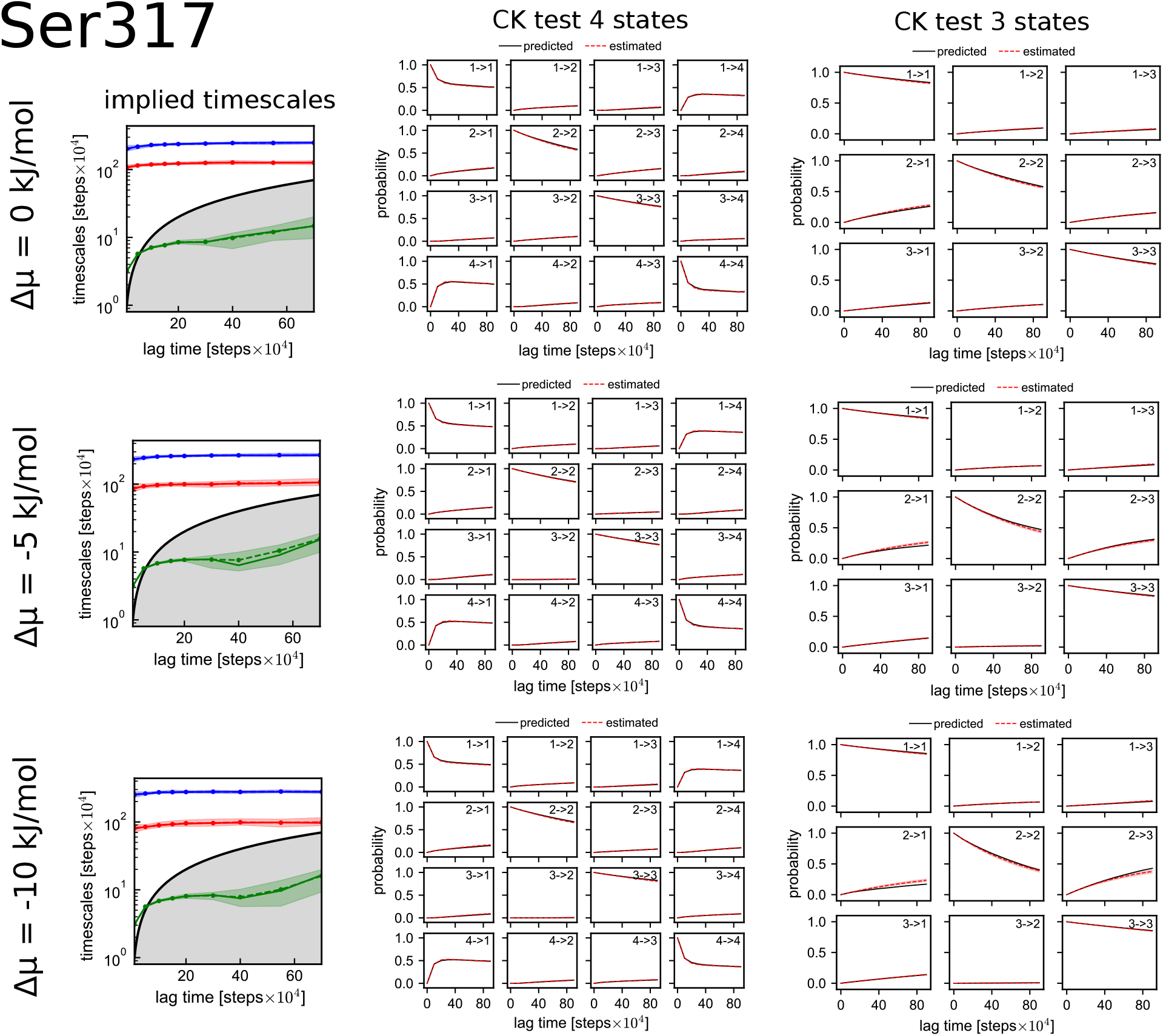
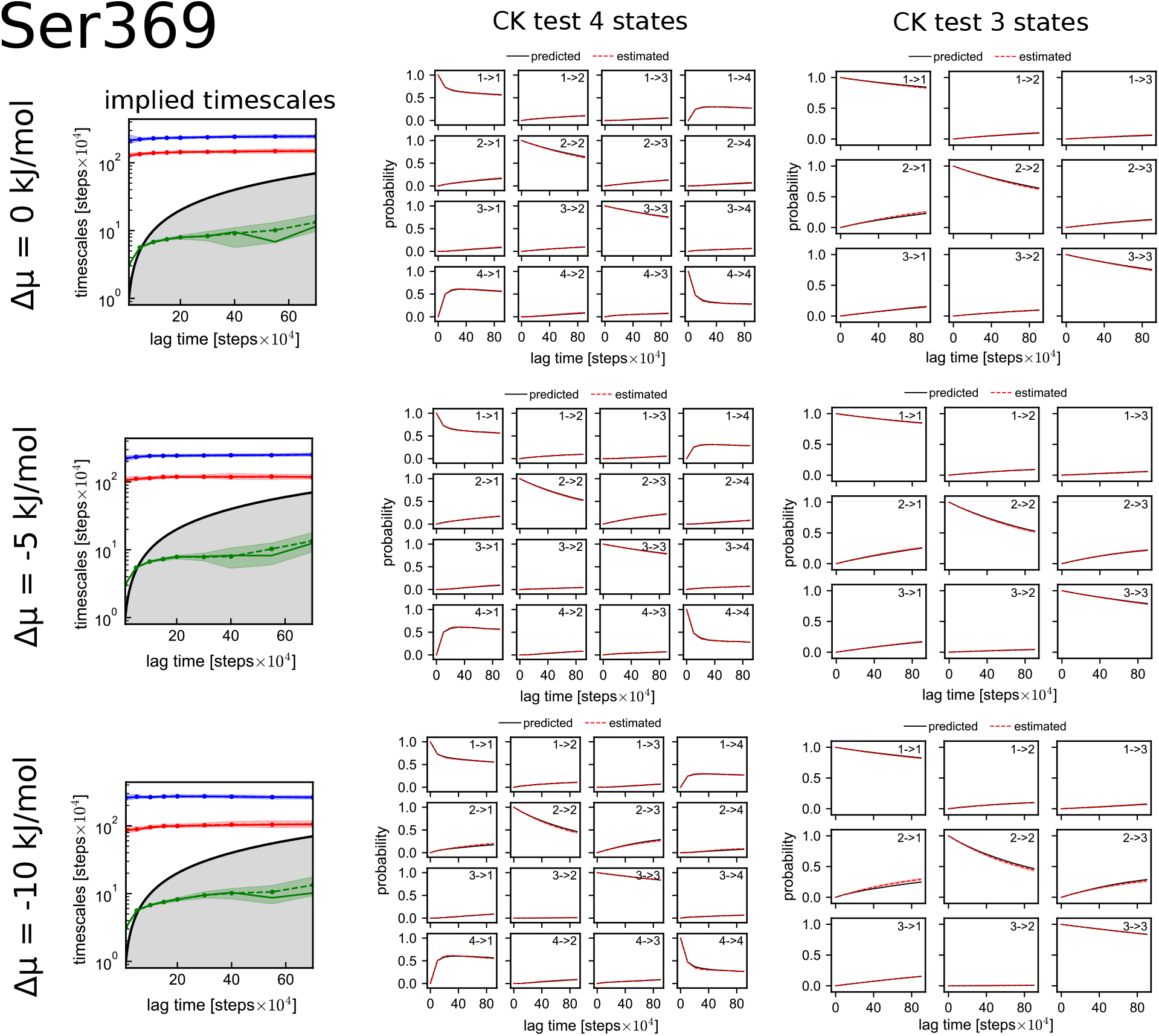
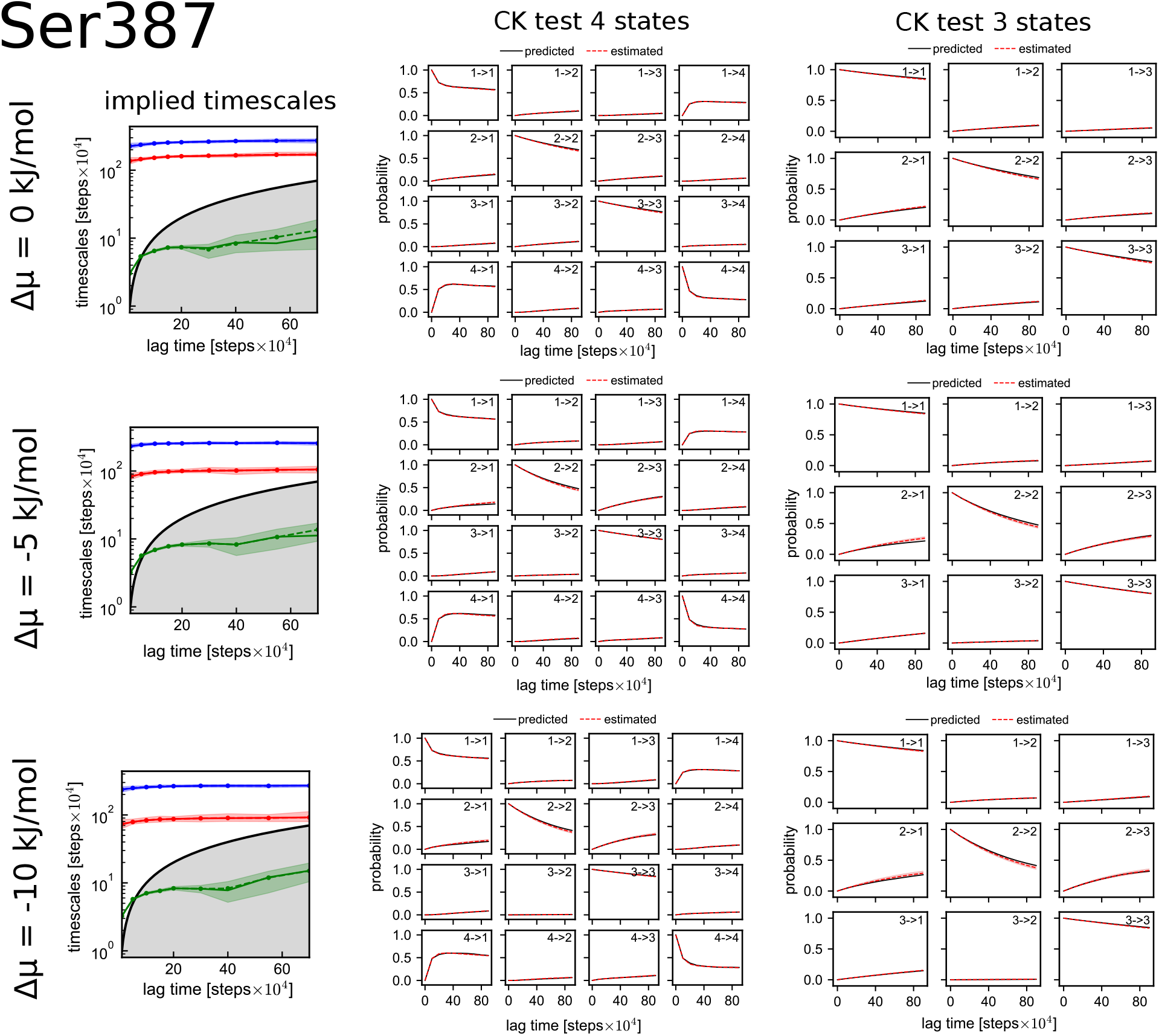
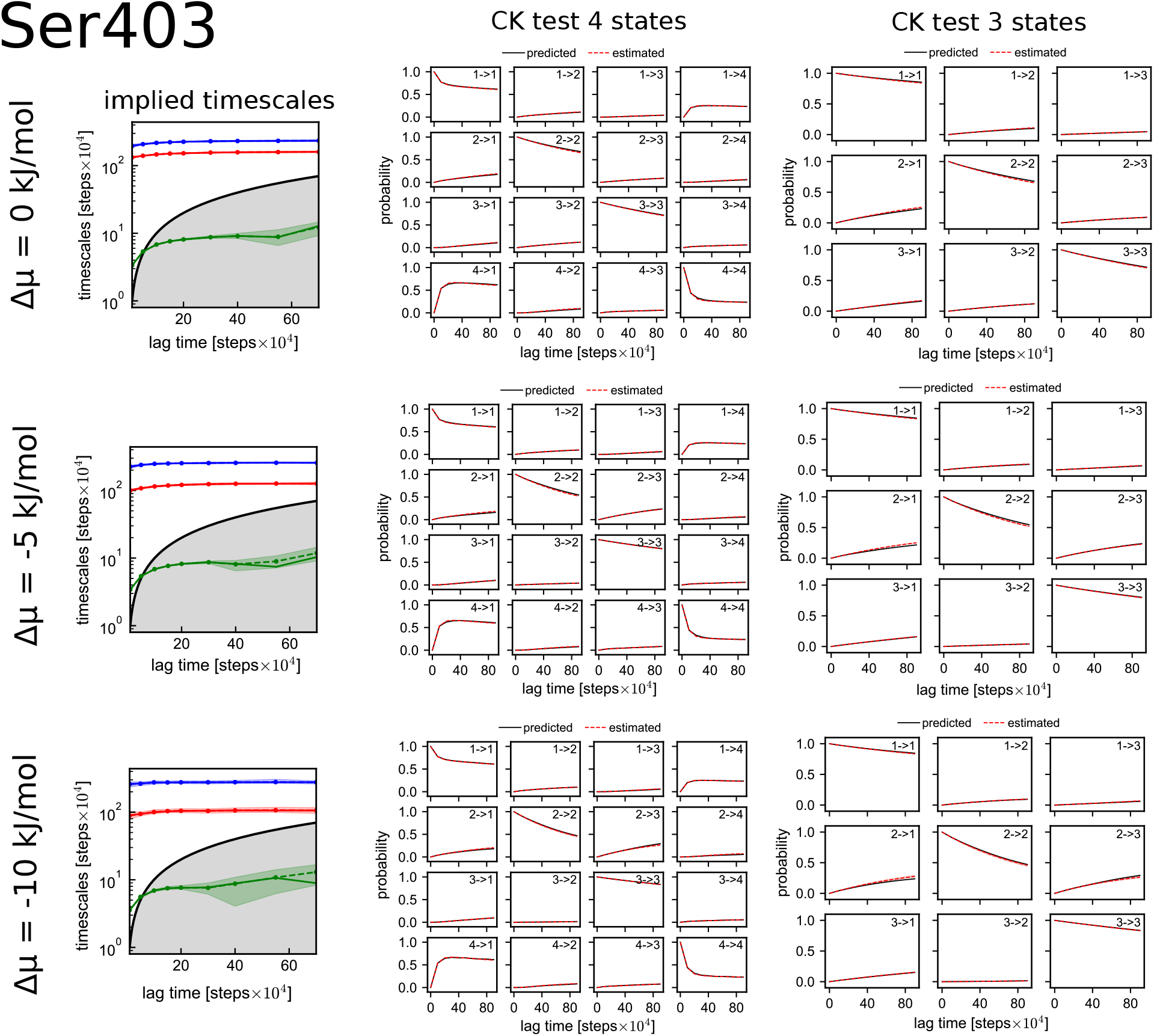
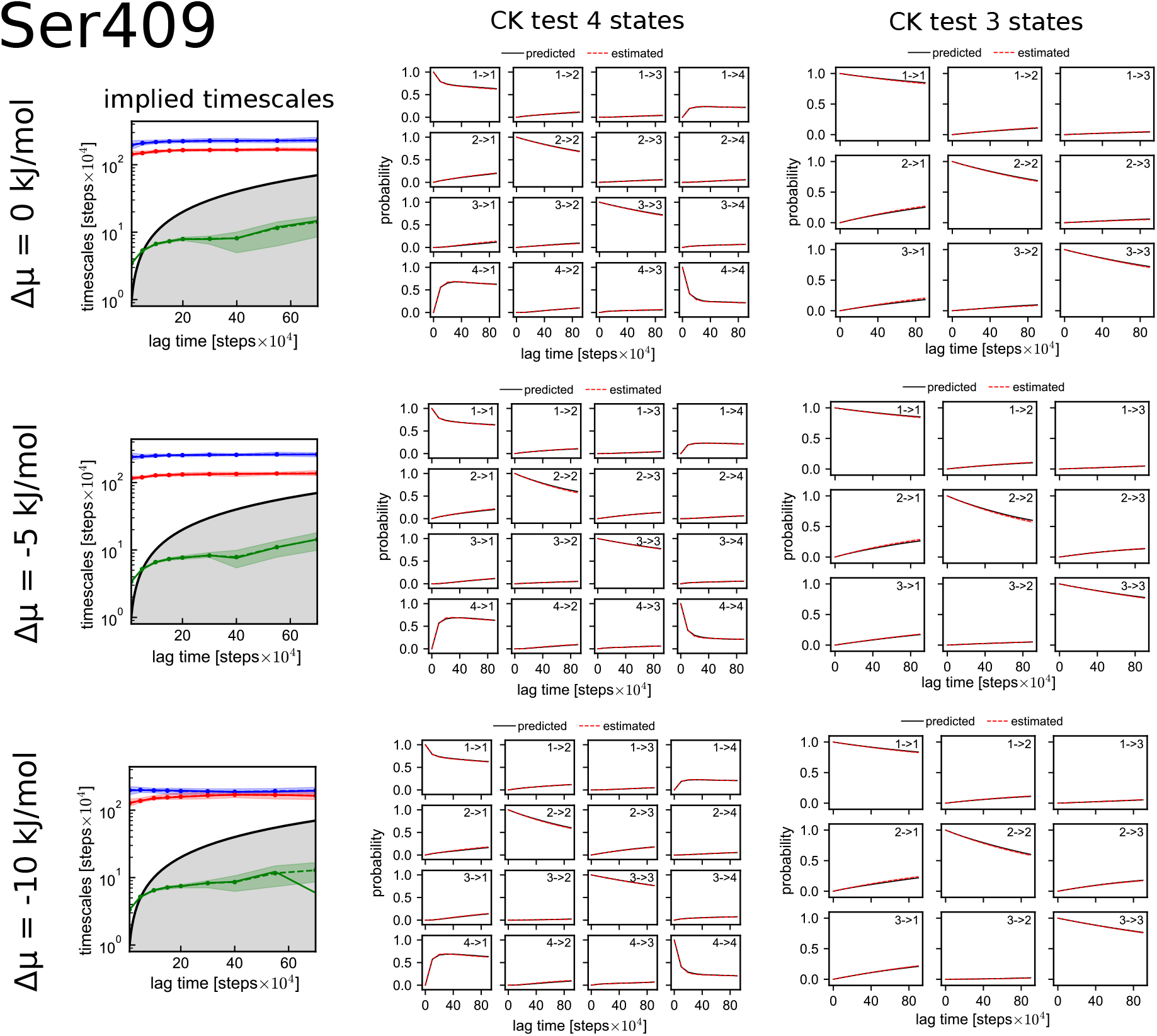
Implied timescales and CK test for every validation simulation. We estimated Δ*µ*_cycle_ also with a 3-state MSM, merging together state 1 and 4 into the new state 1. We report the CK test also for the 3-state MSM case. The implied timescales are constant for *τ* ≥ 10 Markov chain steps. The smaller implied timescale is smaller than the lag time for *τ* ≥10 Markov chain steps. The CK test confirm that the MSM correctly fulfill the markovianity condition.

**Figure S2:**
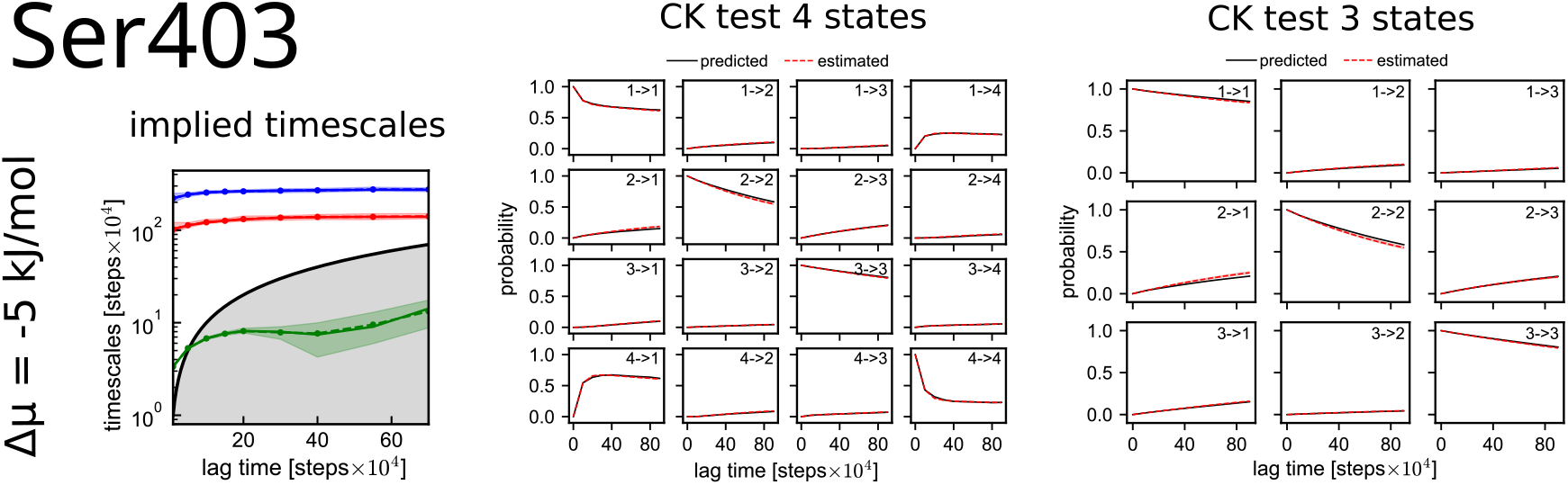
Implied timescales and CK test for Ser403 and Δ*µ*_P_ = − 5 kJ/mol using a version of VAMPnet with 4260 input distances. We estimated Δ*µ*_cycle_ also with a 3-state MSM, merging together state 1 and 4 into the new state 1. We report the CK test also for the 3-state MSM case.

### Phosphorylation modifies interaction of CK1*δ* with TDP-43 LCD

#### Binding free energy

We computed the binding free energy Δ*G*_*bind*_ between 1) CK1*δ* folded-domain and wild type TDP-43 LCD, 2) full-length CK1*δ* and wild type TDP-43 LCD, 3) CK1*δ* folded-domain and triple phosphorylated TDP-43 LCD (pSer 395, pSer 403, pSer 410). For the first 2 cases, we used the data from the equilibrium simulations without phosphorylation step (450 *µ*s for case 1) and 900 *µ*s for case 2)), while for case 3) we collected 20 *µ*s of simulation time.

The binding free energy is estimated as

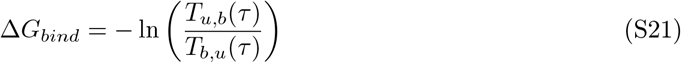

where *T*_*u,b*_(*τ*) is the probability to have a bound state at time *t* + *τ*, given an unbound state at time *t*, and *T*_*b,u*_(*τ*) is the probability to have an unbound state at time *t* + *τ*, given a bound state at time *t*. In order to get the transition probabilities, we discretize the simulation trajectories in bound and unbound state using VAMPnet (in the same way as in Fig. 2, as explained in Methods) and estimate *T*_*u,b*_ and *T*_*b,u*_ at lag time *τ* = 5 × 10^4^ MD steps using a Maximum Likelihood estimator for MSM [50].

#### Plots

**Figure S3:**
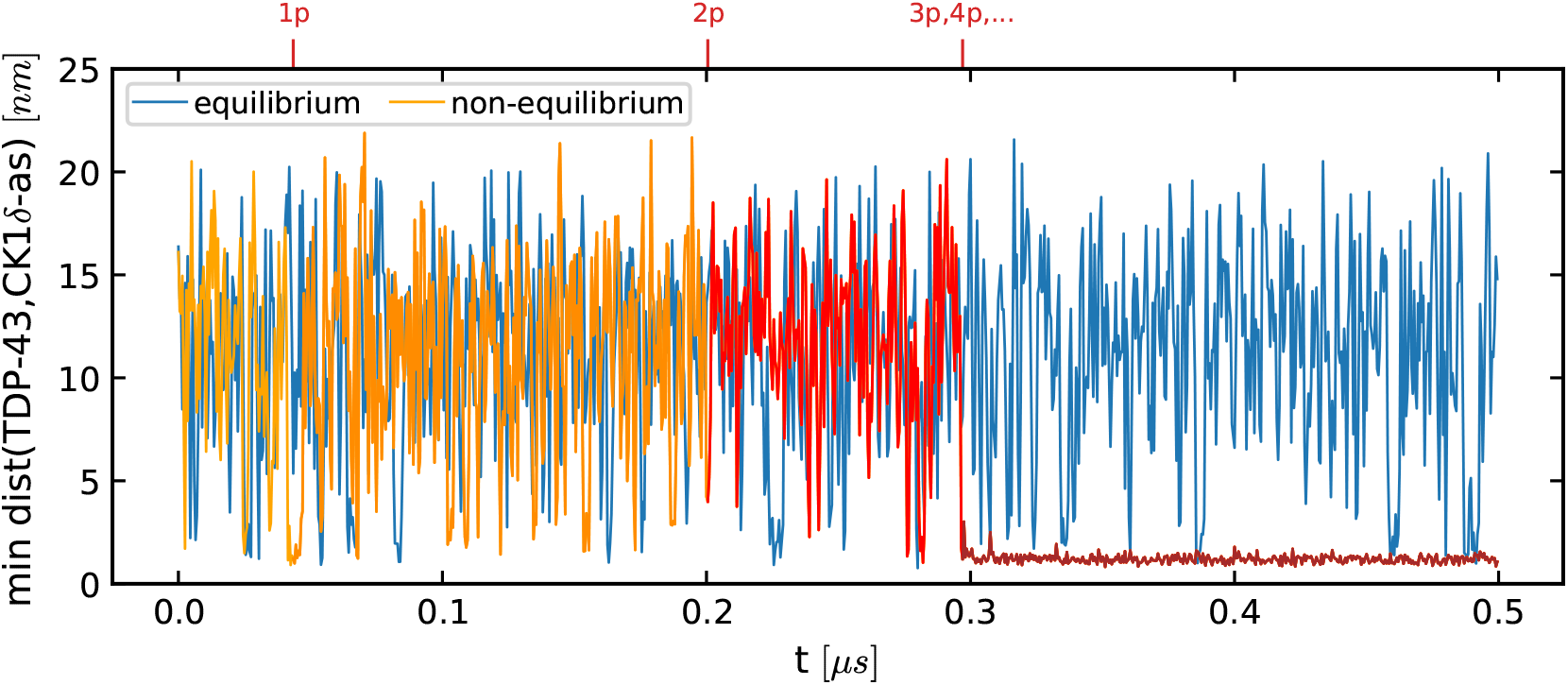
Example trajectory of minimum distance between residues of TDP-43 LCD and the active site of CK1*δ* folded-domain in equilibrium simulation without phosphorylation (blue) and in non-equilibrium simulation (orange) in dilute concentration. The color of the non-equilibrium trajectory becomes darker after every phosphorylation event. In this example TDP-43 stays bound to the enzyme after 3 phosphorylations.

**Figure S4:**
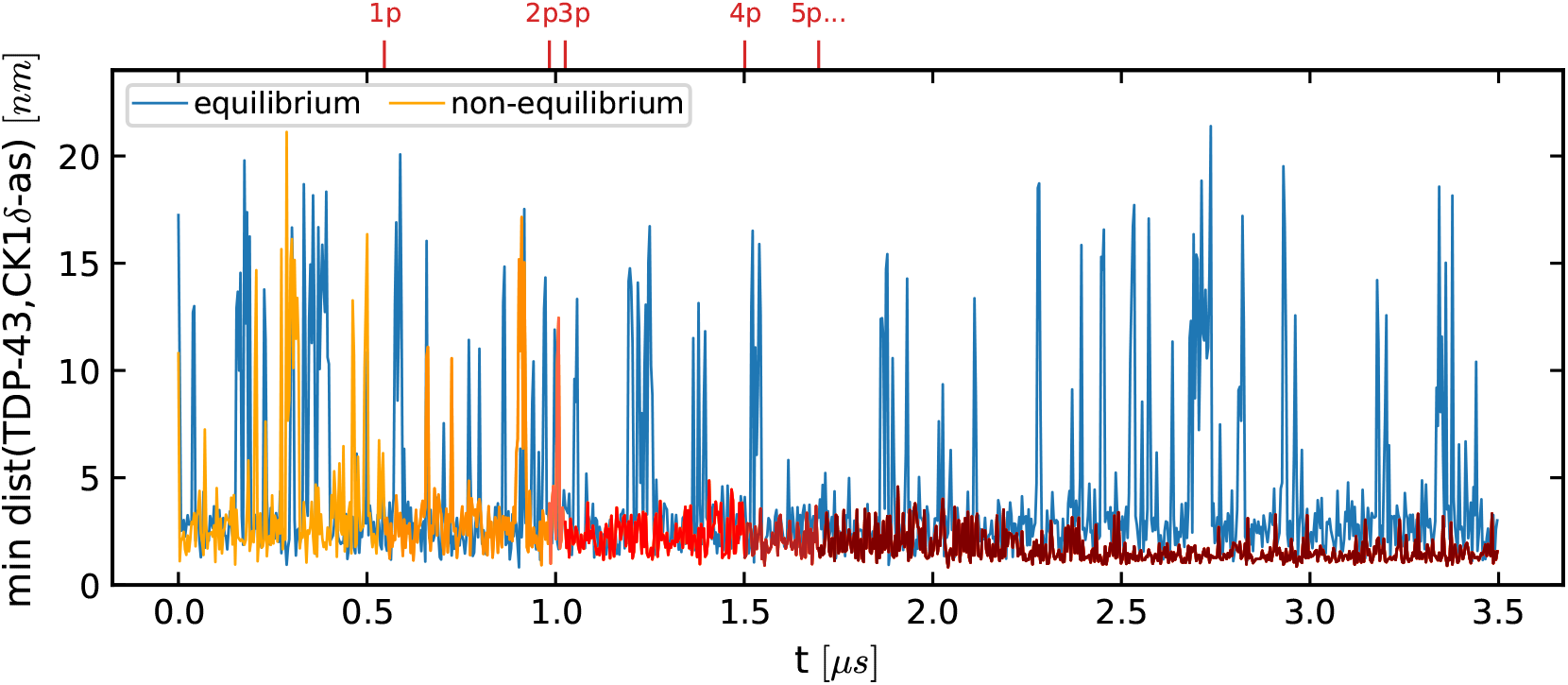
Example trajectory of minimum distance between residues of TDP-43 LCD and the active site of full-length CK1*δ* in equilibrium simulation without phosphorylation (blue) and in non-equilibrium simulation (orange) in dilute concentration. The color of the non-equilibrium trajectory becomes darker after every phosphorylation event. In this example TDP-43 stays bound to the enzyme after 3 phosphorylations.

**Figure S5:**
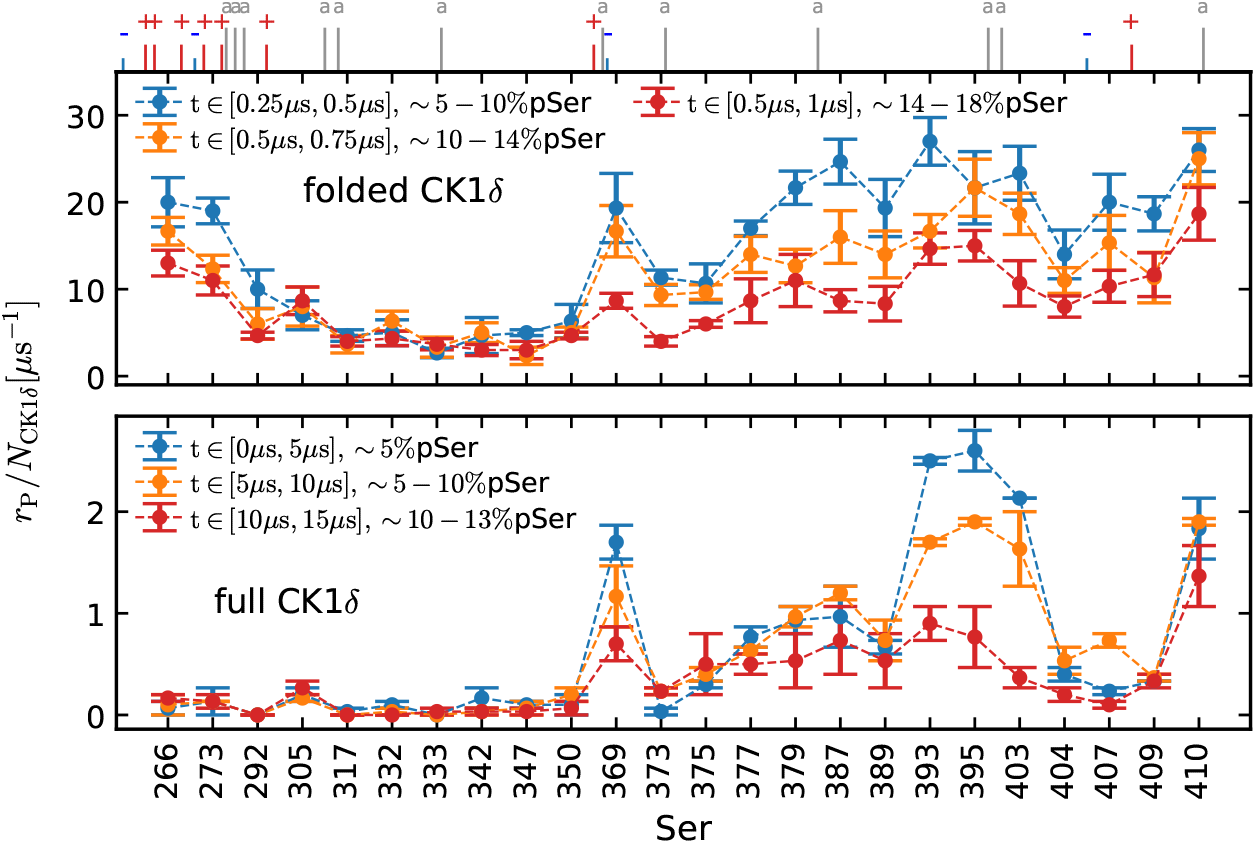
Phosphorylation rates *r*_P_ for every Ser of TDP-43 LCD in presence of 3 CK1*δ* folded-domain (top panel) or 3 full-length CK1*δ* (lower panel) in condensate for different parts of the trajectory. The ticks on top show the position of the charged and aromatic residues. Phosphorylation rates of the most phosphorylated Ser decrease with time.

**Figure S6:**
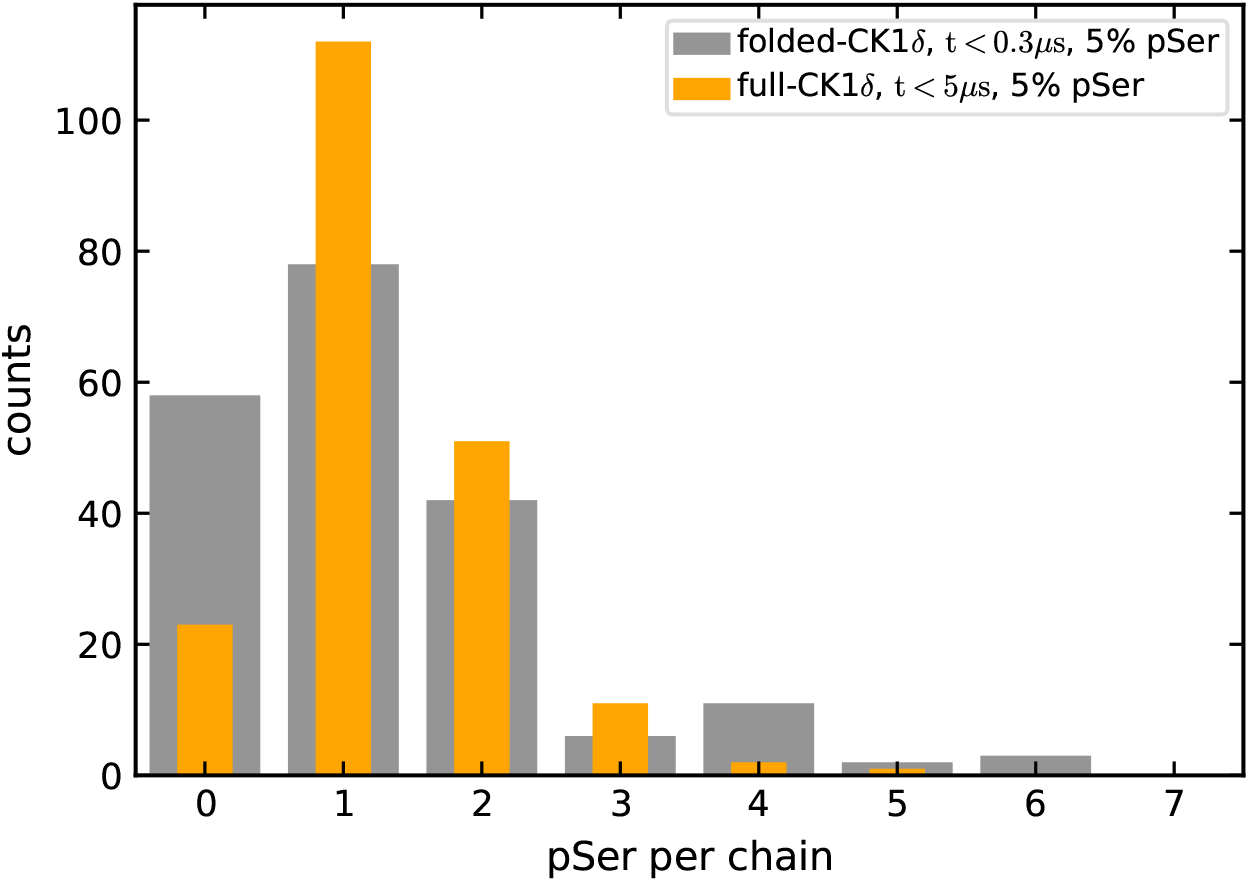
Histogram of the presence of phosphate per chain after 0.3 *µ*s for the simulation of TDP-43 condensate with 3 CK1*δ* folded-domain (grey) and after 5 *µ*s for the simulation of TDP-43 condensate with 3 full-length CK1*δ* (orange). In both cases about 5% of total number of Ser are phosphorylated at the time of the measurement.

### CK1*δ* active sites and charges

**Figure S7:**
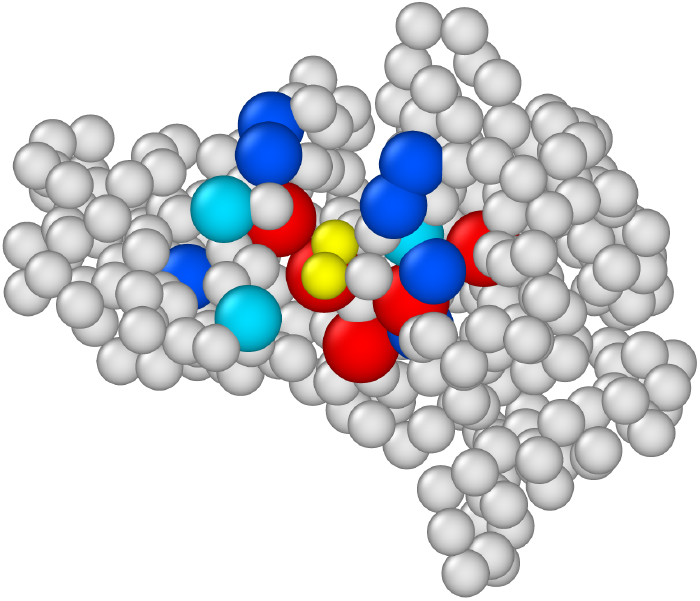
CK1*δ* with colored charged residue close to the active site. In blue the +e charged residues, in light blue His residues (considered +0.5e in our simulations), in red -e charged residues. In yellow the active site residues.

### Phosphorylation acceptance

**Figure S8:**
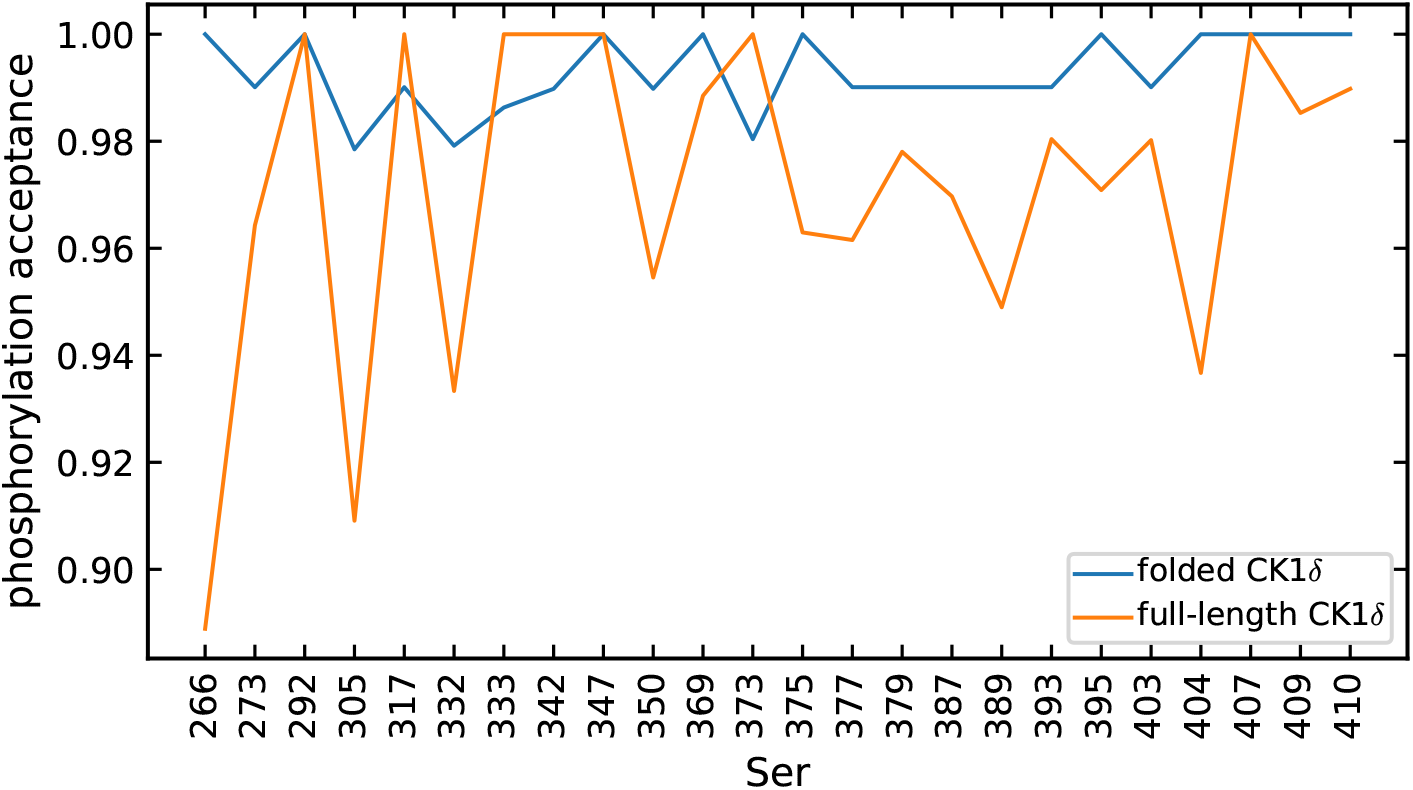
Acceptance ratio for phosphorylation step for simulations with CK1*δ* folded-domain (blue) and full-length CK1*δ* (orange). The acceptance ratio is very close to 100% for every Ser of TDP-43 LCD.

### Phosphorylation process fit

**Figure S9:**
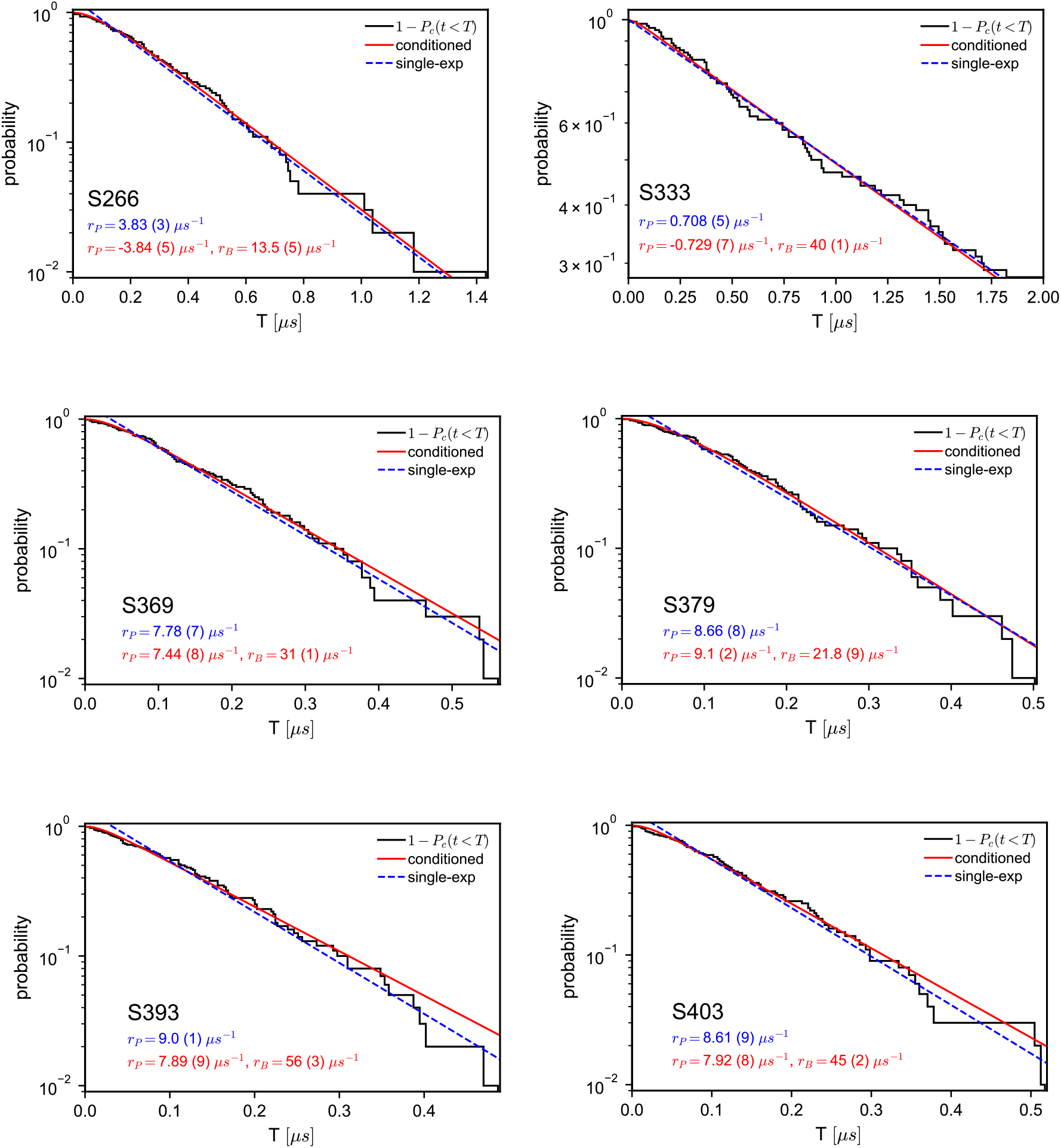
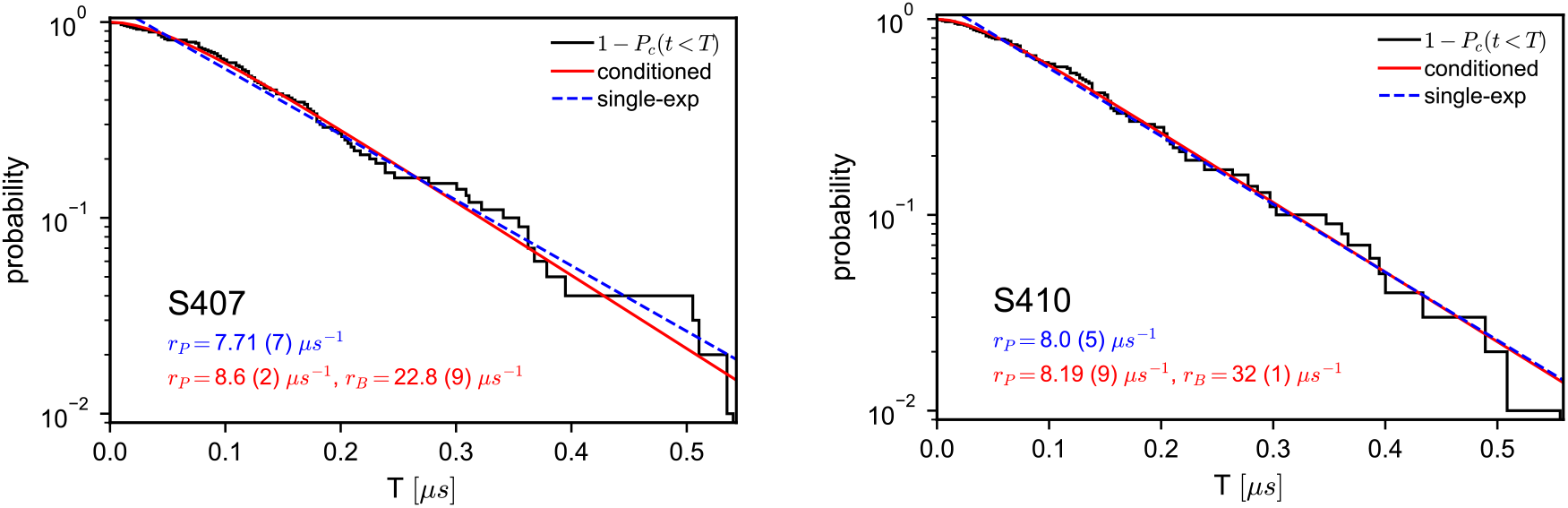
Normalized inverse cumulative histograms of phosphorylation times (black solid lines) and fit with simple single-exponential process (blue dotted lines, rate estimates in blue) and conditioned single-exponential process (red solid lines, rates estimates in red) for 8 different Ser residues. Most of the times the conditioned exponential process fits perfectly. The rate extrapolations from the two fits are in agreement with the bayesian estimates. The rate *r*_*B*_ is different for every serine.

### Phosphorylation rank

**Figure S10:**
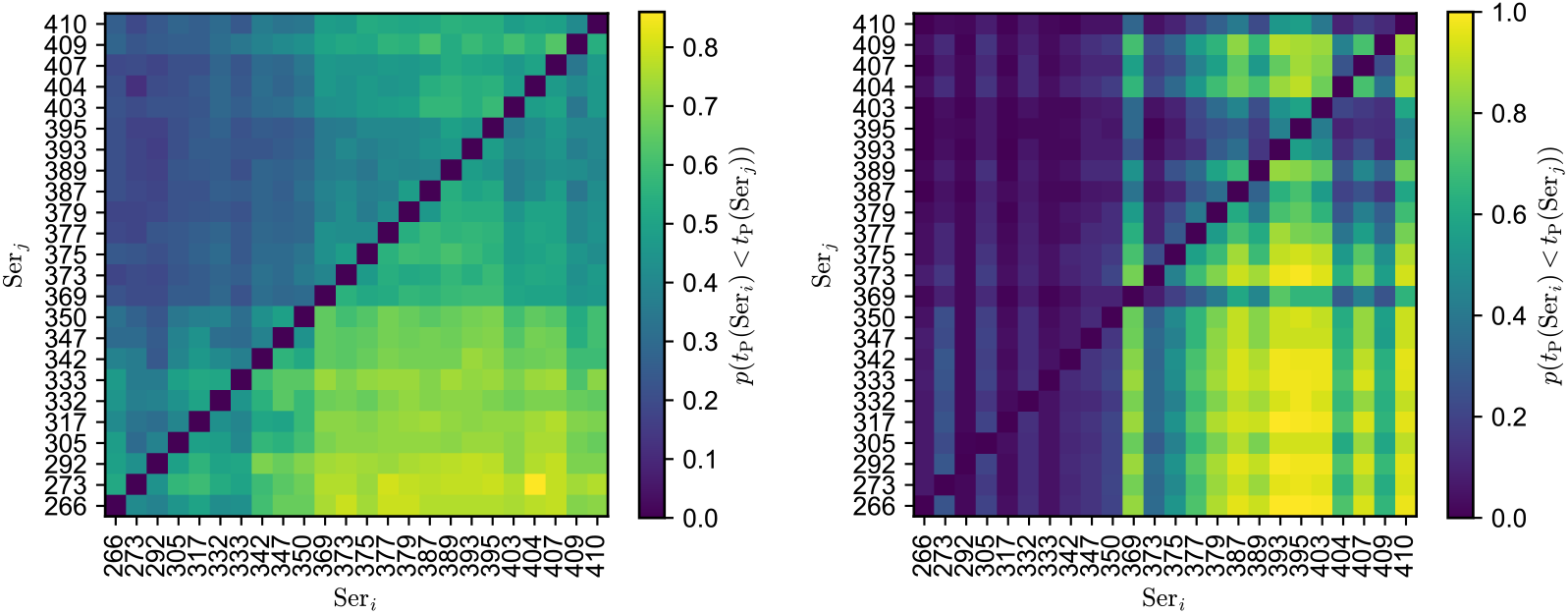
Probability *p*(*t*(Ser_*i*_) *< t*(Ser_*j*_)) of Ser_*i*_ being phosphorylated ahead of Ser_*j*_ for system with averaged-interaction chain and CK1*δ* folded-domain (left) and for the system with TDP-43 LCD wild type and full-length CK1*δ* (right), data from 100 trajectories.

**Figure S11:**
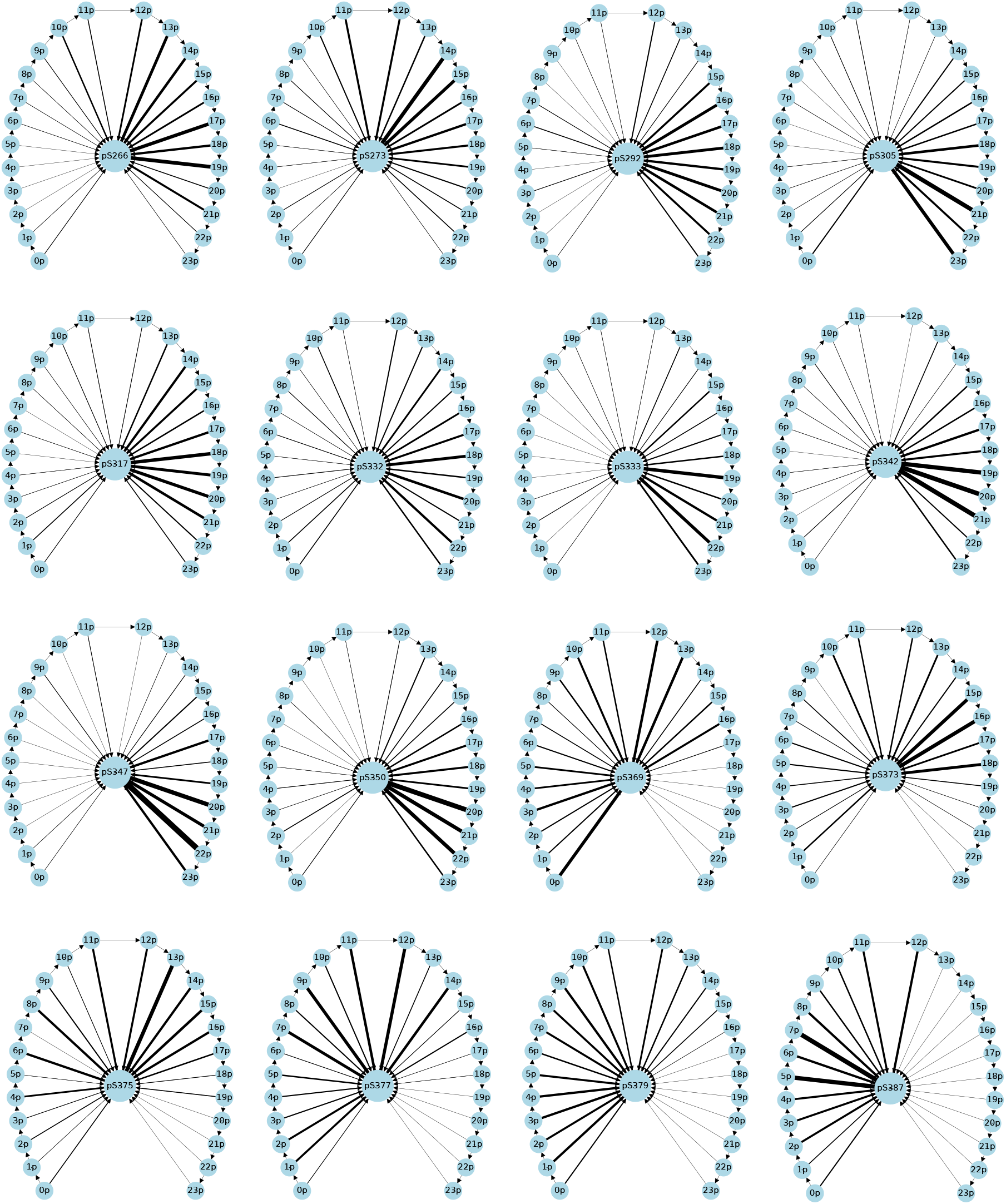
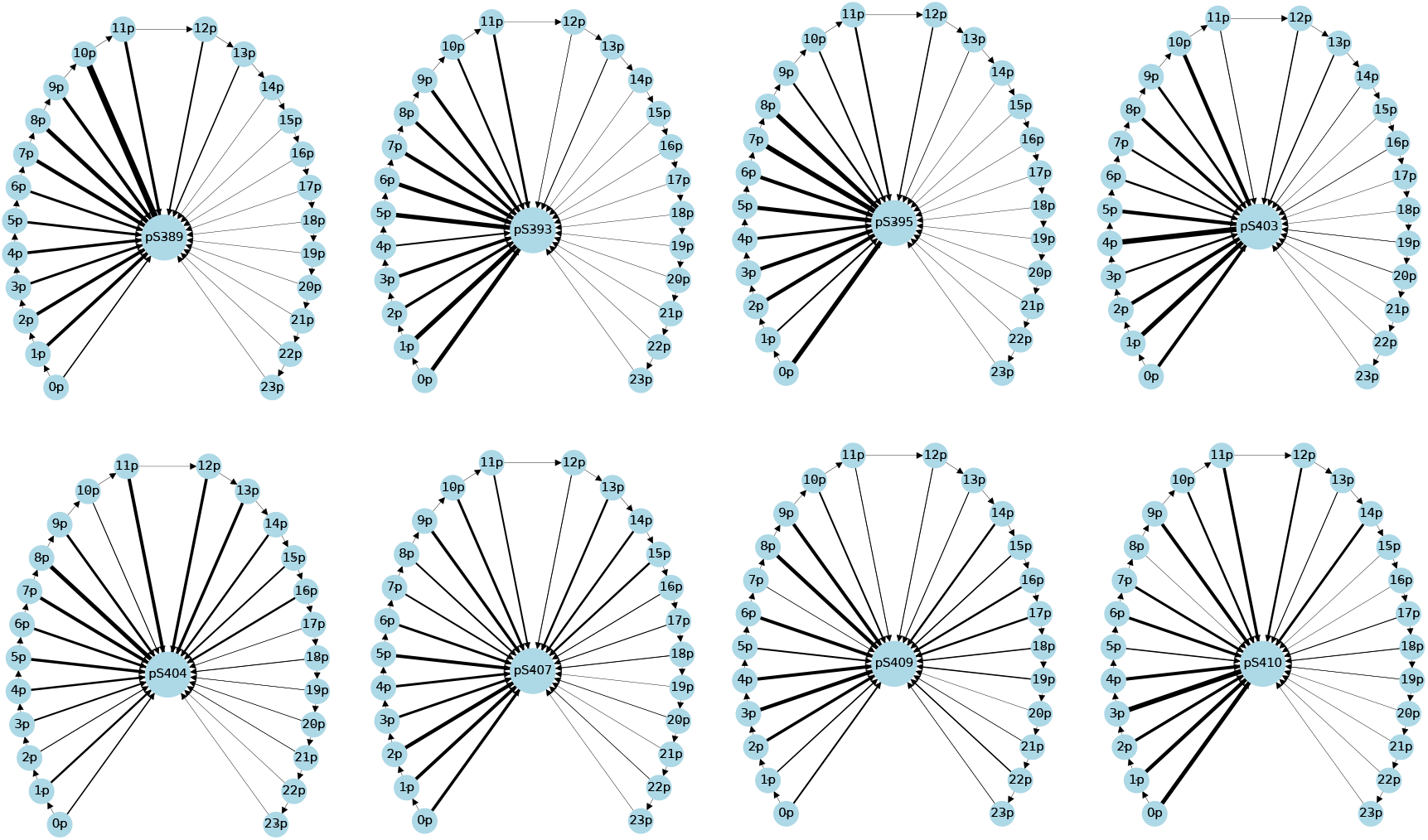
Phosphorylation pattern representation for every Ser of TDP-43 LCD. The thickness of the arrows represent the percentage of simulations in which the Ser in the center of the graph was phosphorylated after n other Ser residues.

#### Correlation plots: *r*_P_ vs *r*_c_ in dilute regime for full-length CK1*δ*

**Figure S12:**
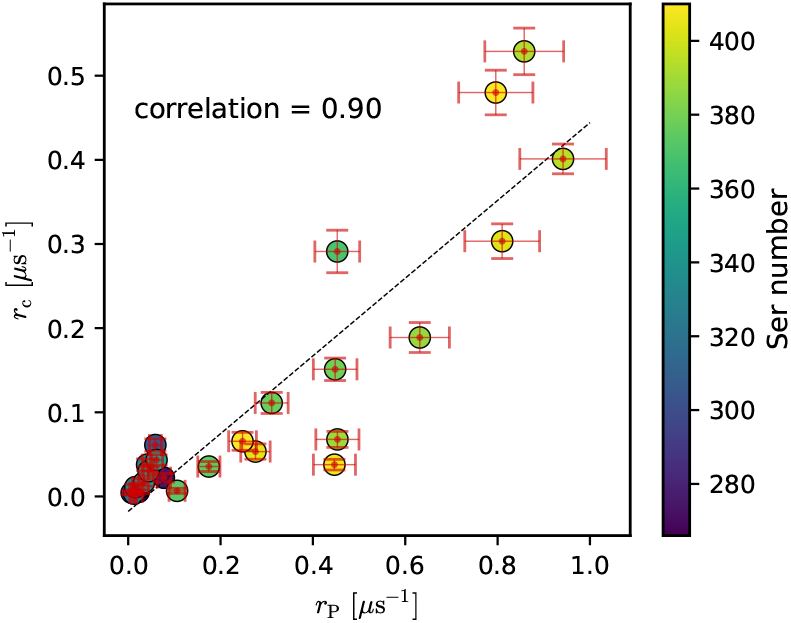
Correlation plots of contact frequency in equilibrium *r*_c_ and phosphorylation rates *r*_P_ in dilute regime for simulations with full-length CK1*δ*.

#### Correlation plots: dilute vs dense regime phosphorylation rates

**Figure S13:**
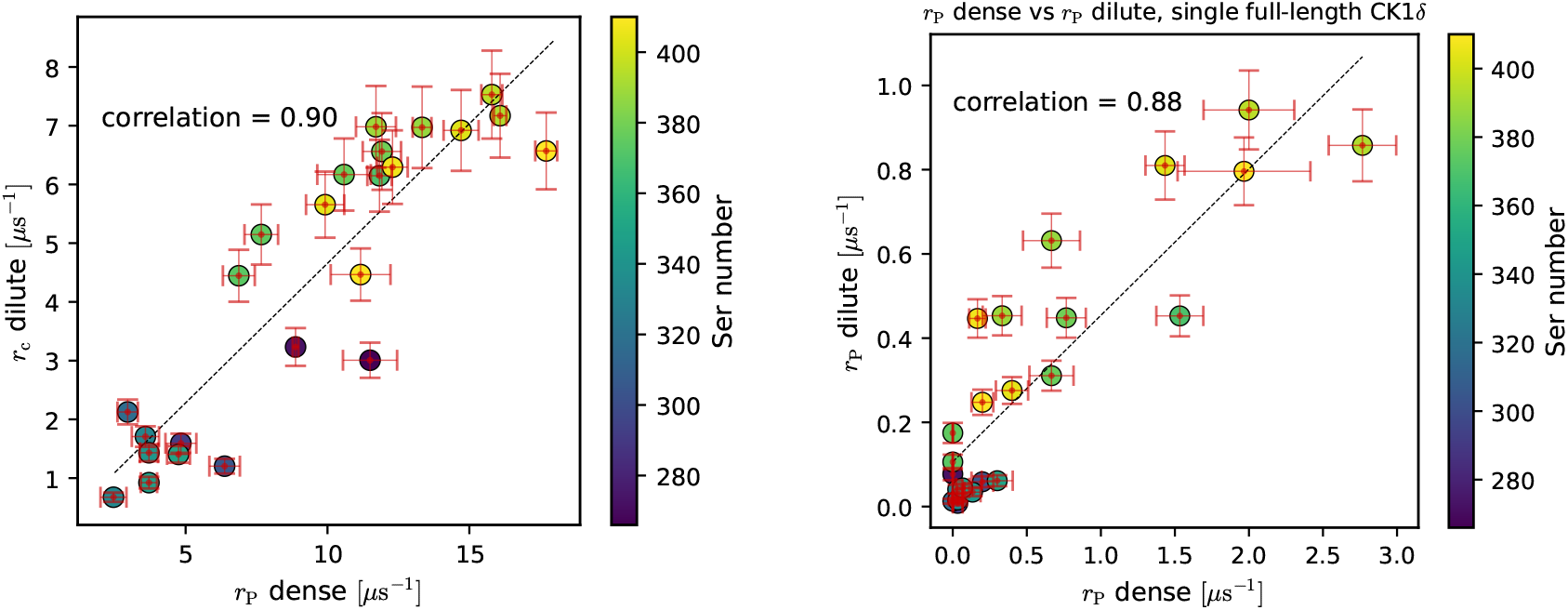
Correlation plots of phosphorylation rates *r*_P_ in dense (x-axis) and dilute (y-axis) regimes for simulations with one CK1*δ* folded-domain (left) and one full-length CK1*δ* (right).

#### Correlation plots: *r*_P_ vs *r*_c_ in condensate

**Figure S14:**
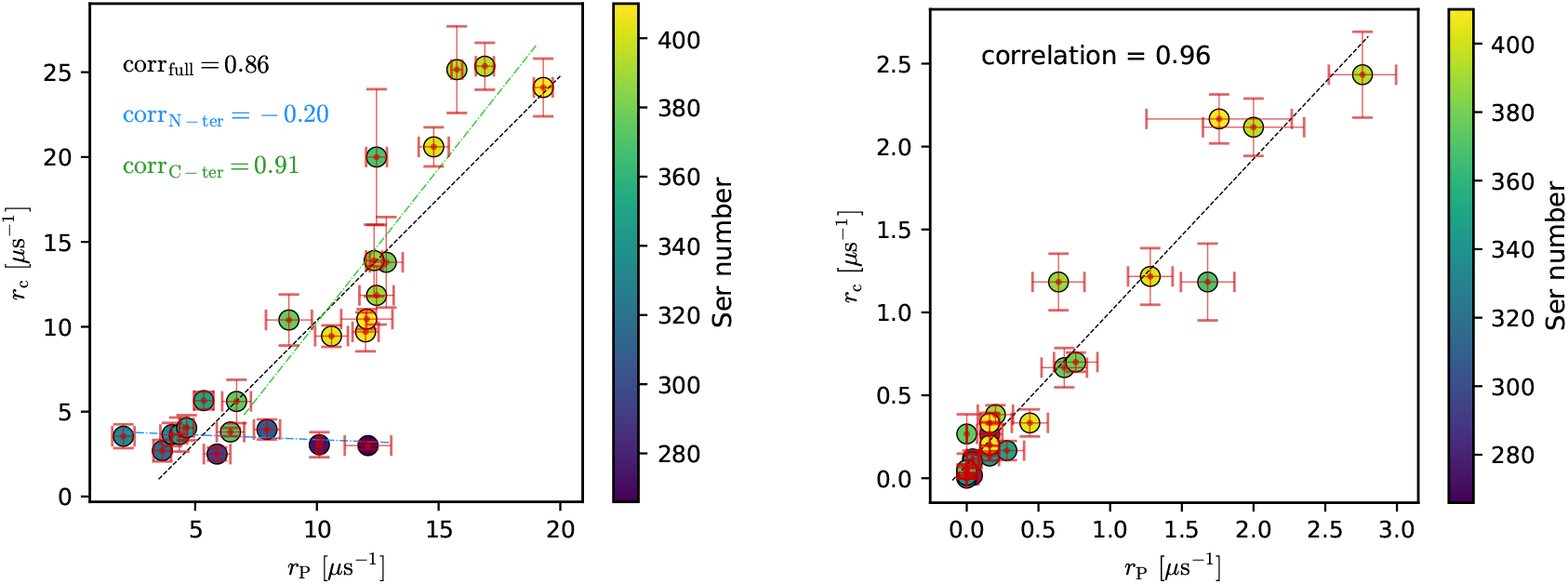
Correlation plots of contact frequency in equilibrium *r*_c_ and phosphorylation rates *r*_P_ in condensate for simulations with CK1*δ* folded-domain (left) and full-length CK1*δ* (right).

#### CK1*δ* disordered domain do not cover the active site

**Figure S15:**
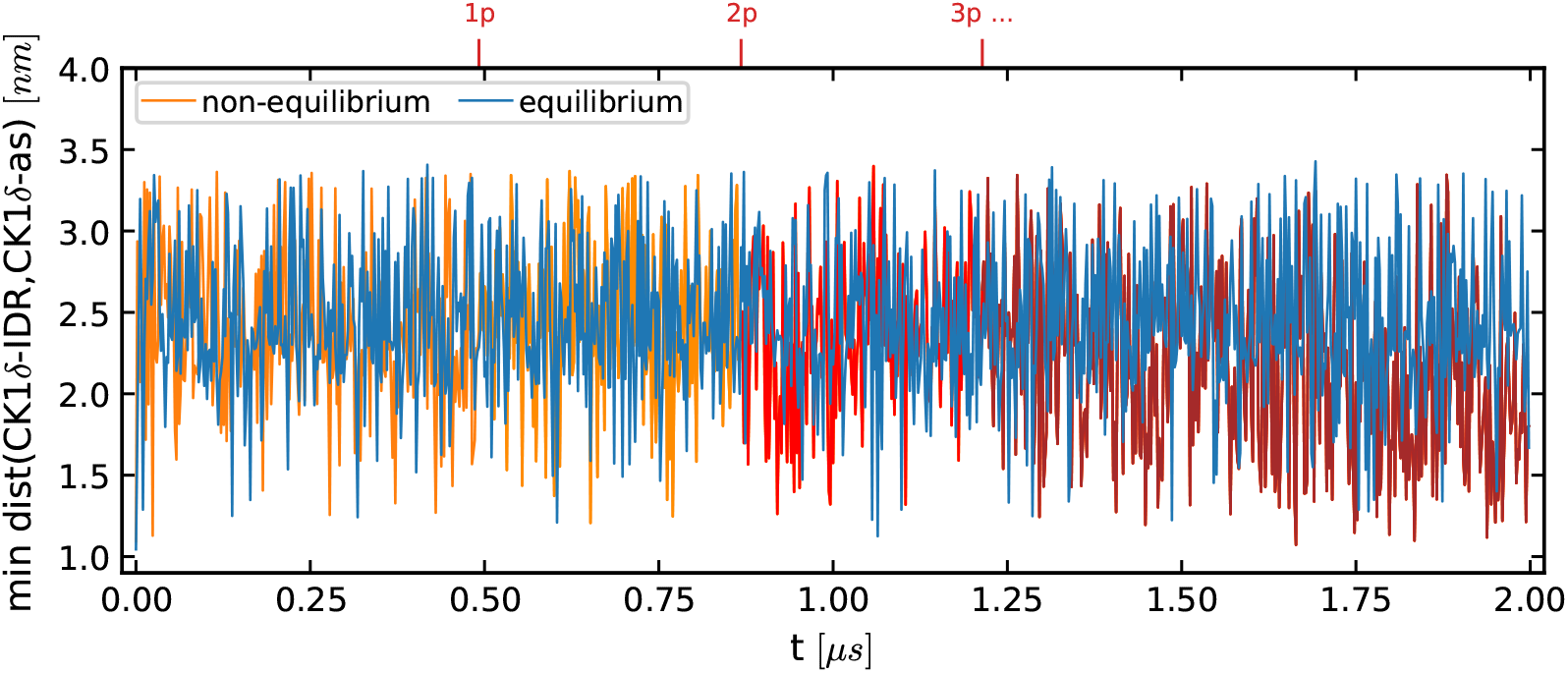
Example trajectory of minimum distance between residues of full-length CK1*δ* IDR and its active site in equilibrium simulation without phosphorylation (blue) and in non-equilibrium simulation (orange) in dilute concentration. The color of the non-equilibrium trajectory becomes darker after every phosphorylation event. The distance never stays stable around the contact distance, but oscillates around larger values, suggesting that the CK1*δ* IDR does not cover the active site for an extended time interval.

**Figure S16:**
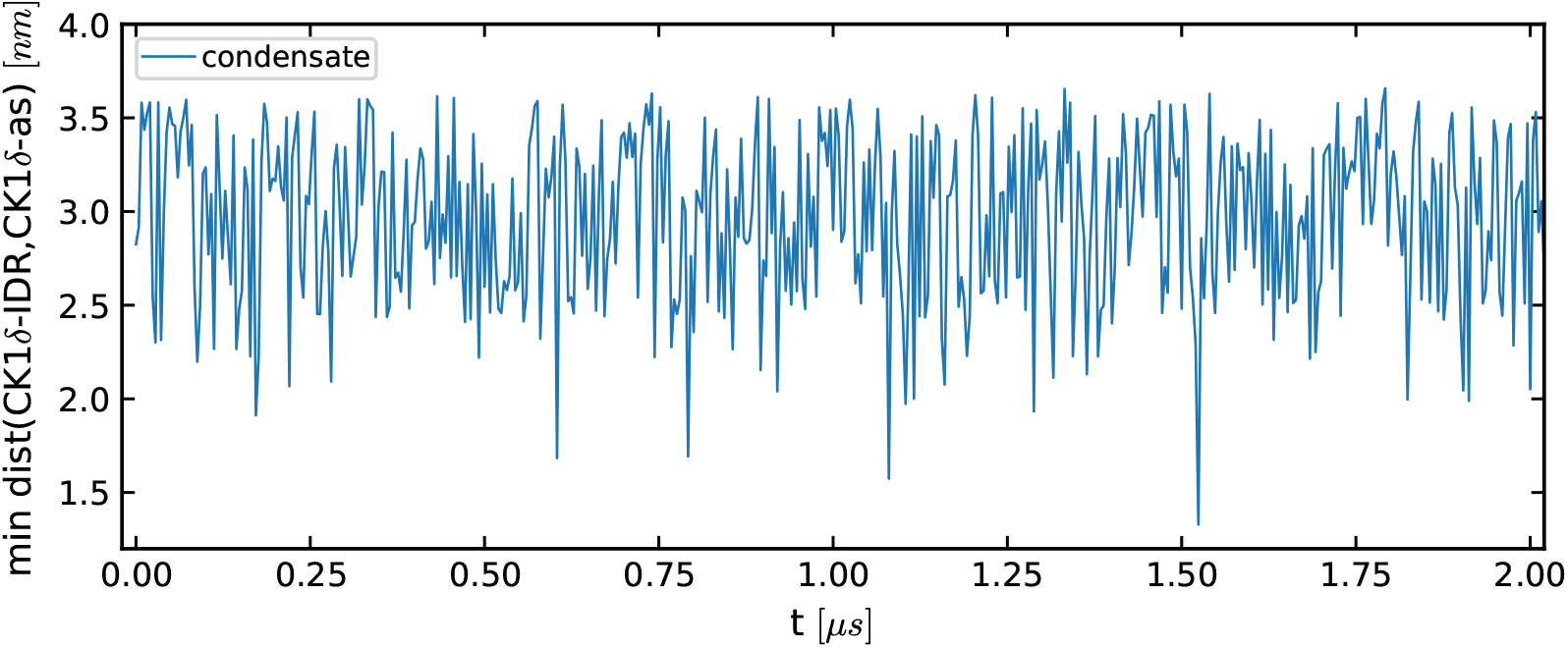
Example trajectory of minimum distance between residues of full-length CK1*δ* IDR and its active site in condensate simulations. The distance never stays stable around the contact distance, but oscillates around larger values, suggesting that the CK1*δ* IDR does not cover the active site for an extended time interval.

### Movies

**SI Movie 1**

Movie from simulation of single TDP-43 LCD chain and single CK1*δ* folded domain with phosphorylation step (Eq. 2) and reservoir exchange step (Eq. 4) in cubic box of 50 nm side length using HPS model. In this simulation, the only phosphosite is Ser403 and Δ*µ*_P_ = − 5 kJ/mol. This simulation was used in Fig. 2.

**SI Movie 2**

Movie from simulation of single TDP-43 LCD chain and single CK1*δ* folded domain with phosphorylation step in cubic box of 30 nm side length using modified HPS model. This simulation was used in Fig. 3e.

**SI Movie 3**

Movie from simulation of 200 TDP-43 LCD chains and 3 CK1*δ* folded-domain with phosphorylation step in cubic box of 100 nm side length using modified HPS model.

**SI Movie 4**

Movie from simulation of single TDP-43 LCD chain and single full-length CK1*δ* with phosphorylation step in cubic box of 30 nm side length using modified HPS model.

**SI Movie 5**

Movie from simulation of 200 TDP-43 LCD chains and 3 full-length CK1*δ* with phosphorylation step in cubic box of 100 nm side length using modified HPS model.

